# The brain selectively allocates energy to functional brain networks under cognitive control

**DOI:** 10.1101/2024.12.06.627178

**Authors:** Majid Saberi, Jenny R. Rieck, Shamim Golafshan, Cheryl L. Grady, Bratislav Misic, Benjamin T. Dunkley, Ali Khatibi

**Affiliations:** Neurosciences & Mental Health Program, The Hospital for Sick Children Research Institute, Toronto, Canada; Headache and Orofacial Pain Effort (H.O.P.E.) Laboratory, Department of Biologic and Materials Sciences & Prosthodontics, University of Michigan School of Dentistry, Ann Arbor, Michigan, USA; Rotman Research Institute, Baycrest Health Sciences, Toronto, M6A 2E1, Canada; Department of Psychology, Florida International University, Miami, FL, USA; Psychology, University of Toronto, 100 St. George Street, Toronto, ON, M5S 3G3, Canada; Psychiatry, University of Toronto, Toronto, M5T 1R8, Canada; Department of Neurology and Neurosurgery, McGill University, Montreal, QC, Canada; McConnell Brain Imaging Centre, Montreal Neurological Institute and Hospital, Montreal, QC, Canada; Department of Diagnostic Imaging, The Hospital for Sick Children, Toronto, Canada; Department of Medical Imaging, University of Toronto, Toronto, Canada; Centre of Precision Rehabilitation for Spinal Pain (CPR Spine), School of Sport, Exercise and Rehabilitation Sciences, University of Birmingham, Birmingham, UK; Centre for Human Brain Health, University of Birmingham, Birmingham, UK; Institute for Mental Health, University of Birmingham, Birmingham, UK

**Keywords:** Network energy, cognitive control, executive functions, functional connectivity, canonical functional networks, structural balance theory, brain biomarker, predictive modeling

## Abstract

Network energy has been conceptualized based on structural balance theory in the physics of complex networks. We utilized this framework to assess the energy of functional brain networks under cognitive control and to understand how energy is allocated across canonical functional networks during various cognitive control tasks. We extracted network energy from functional connectivity patterns of subjects who underwent fMRI scans during cognitive tasks involving working memory, inhibitory control, and cognitive flexibility, in addition to task-free scans. We found that the energy of the whole-brain network increases when exposed to cognitive control tasks compared to the task-free resting state, which serves as a reference point. The brain selectively allocates this elevated energy to canonical functional networks; sensory networks receive more energy to support flexibility for processing sensory stimuli, while cognitive networks relevant to the task, functioning efficiently, require less energy. Furthermore, employing network energy, as a global network measure, improves the performance of predictive modeling, particularly in classifying cognitive control tasks and predicting chronological age. Our results highlight the robustness of this framework and the utility of network energy in understanding brain and cognitive mechanisms, including its promising potential as a biomarker for mental conditions and neurological disorders.

## Introduction

Over recent years, the perspective of brain energy modeling has increasingly captivated the attention of neuroimaging scientists. This approach, rooted in the principles of statistical physics, offers a powerful framework for examining the collective behavior of brain components and their interactions. By focusing on the brain’s optimization strategies, it provides an energy landscape that identifies potential brain states and highlights the brain’s ability to transition between these states. Tracking brain dynamics through this lens allows researchers to identify critical neural states and understand how the brain manages the opposing demands of maintaining stability while allowing for necessary adaptability.

Numerous studies have explored the brain’s energy landscape of neurophysiological activity using models derived from pairwise maximum entropy [1]. For instance, Gu et al. applied an energy function based on regional activation and interactions, demonstrating that the most probable brain states, corresponding to minimal energy, exhibit consistent activation patterns across brain regions [2]. Similarly, Watanabe et al. described resting-state activity, including the default-mode and frontoparietal regions, as attractor dynamics, where brain states tend to move toward stable energy minima. Their findings revealed that a small number of local energy minima form the backbone of each super-regions, with the majority of states associated with these attractors [3]. In another study, they explored bistable visual perception and showed that brain activity fluctuates between three spatially distributed energy minima: visual-area-dominant, frontal-area-dominant, and intermediate states. They further demonstrated that participants with brain activity patterns favoring the visual-area-dominant state exhibited more stable perception [4]. In a related study, Ashourvan et al. generated an energy landscape where local minima represented attractor states associated with specific patterns of modular structure in the brain. They identified distinct functional communities, characterizing visual, attention, sensorimotor, and subcortical regions by a single community, while linking other regions to executive control or default mode and salience communities [5]. Several other studies have utilized this energy definition approach to investigate various aspects of brain dynamics and function, highlighting its applicability across different neural conditions and disorders [6-9].

By focusing on the organization of functional connections from a network modeling viewpoint, structural balance theory has recently been applied to model the energy of complex brain networks [10]. This methodology, initially introduced by Fritz Heider in the context of social science [11-12] and later formalized in the physics of complex networks [13-14], distinguishes between stable and unstable network elements by considering both positive and negative characteristics of connections. Structural balance theory provides a robust framework for investigating the energy landscape of the network by utilizing the occurrence patterns of both stable and unstable elements [13], enabling the monitoring of network dynamics [15-16], transitions between various network states [17-18], and the identification of basins within a network system [19].

The assessment of brain network energy through the lens of structural balance theory provides valuable insights into the organization of functional connections (Figure 1). A lower energy level reflects stability, characterized by a well-coordinated configuration of functional connections that reduces internal conflicts and promotes efficient neural processing, as seen in resting-state conditions. In contrast, a high energy level, indicative of instability, demonstrates tension between brain components and a conflicting configuration of functional connections. It allows dynamic reconfiguration of functional connections, enabling the brain systems to adapt to external demands. These energy dynamics can illustrate the brain’s capacity to shift between stability and instability, essential for managing the competing demands of efficiency and flexibility. Notably, deviations in network energy levels might be associated with disruptions in neural processing, potentially underlying mental health disorders, cognitive impairments, or the effects of therapeutic interventions. In this context, investigating network energy offers key insights into the functional organization of neural systems, with potential applications in enhancing our understanding of brain dynamics and cognitive processes, diagnosing neurological conditions, and optimizing therapeutic interventions.

**Figure 1:**
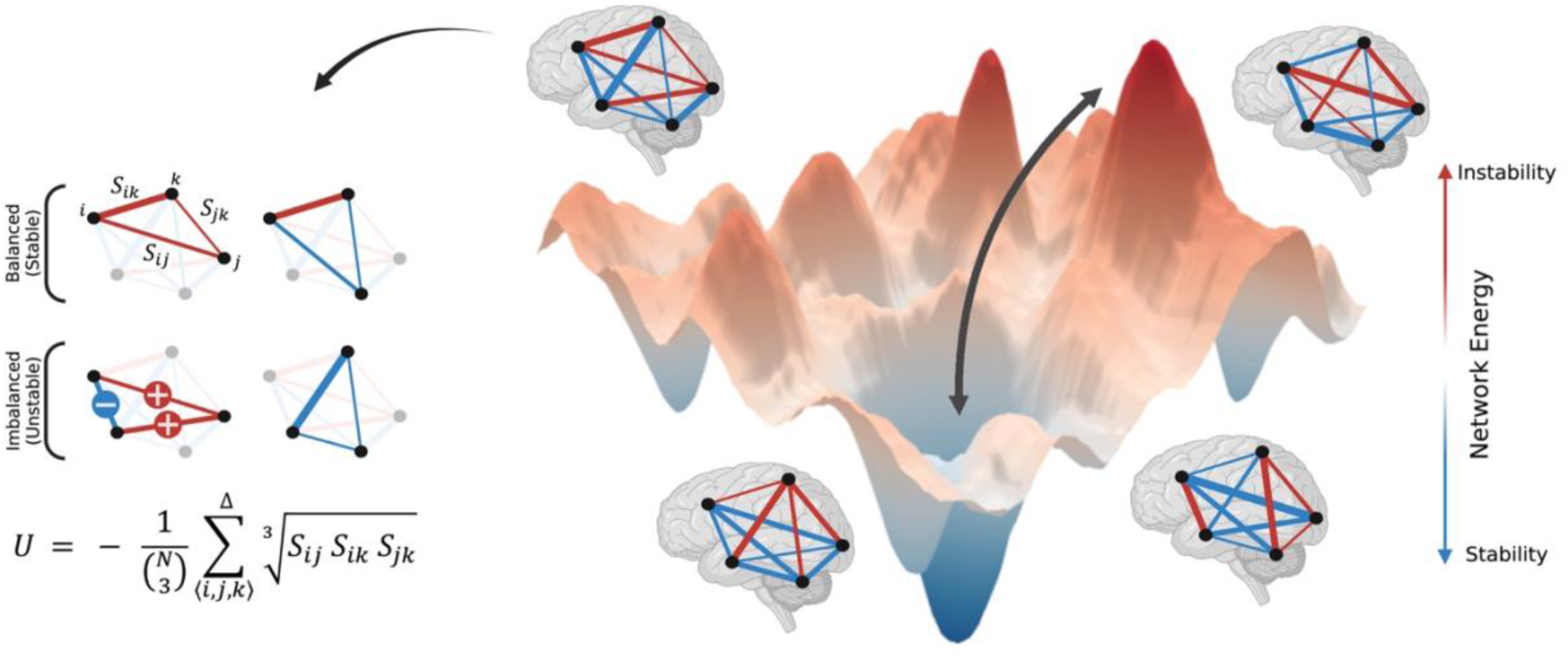
Schematic representation of the brain network’s energy landscape, network states, and state transitions, as well as the calculation of network energy. Brain functional network exhibits different levels of energy, corresponding to the configuration of functional connections. The arrangement of positive (red) and negative (blue) functional connections influences the stability or instability and the level of energy. Triangles formed by functional connections between three brain regions are categorized as either balanced (stable) or imbalanced (unstable), depending on the positive or negative nature of their connections. The intensity of balanced and imbalanced triangles determines the energy level. Network energy is calculated by multiplying the connection weights for each triangle and summing across all triangles in the functional network. Blue and red lines represent negative and positive functional connections, respectively, with the line width indicating the strength of coactivation between two regions. Further details about the formula are provided in the text. (Created with BioRender.com)

Central to this approach is the consideration of the sign and weight of functional connections, which accounts for the interdependence between brain regions. Accordingly, we define the energy of a functional network as:

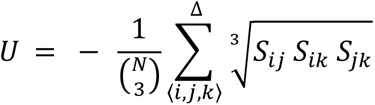

Where *S*_*ij*_ represents the functional connection weight between regions *i* and *j*, assuming either a positive or negative continuous value. The summation is executed across all possible triangles formed by regions *i, j*, and *k*, as indicated by the triangle symbol above the summation. *S*_*ij*_ *S*_*ik*_ *S*_*jk*_ is the multiplication of the triangle’s weighted connections. Positive *S*_*ij*_ *S*_*ik*_ *S*_*jk*_ values denote triangle’s balance, whereas negative *S*_*ij*_ *S*_*ik*_ *S*_*jk*_ values signify triangle’s imbalance. The magnetite of *S*_*ij*_ *S*_*ik*_ *S*_*jk*_ denotes intensity of these effects. The concepts of balance and imbalance are rooted in Fritz Heider’s initial definitions of social interactions between three entities [11-12]; "balance" when a friend’s friend and a foe’s foe are friends, and "imbalance" when a friend’s friend or a foe’s foe become foes. In this context, balanced triangles promote network stability, while imbalanced triangles lead to network instability. The entities within an imbalanced triangle must resolve dissonance to achieve balance over time. As *S*_*ij*_ *S*_*ik*_ *S*_*jk*_ values are positive for balanced triangles and negative for imbalanced triangles, summing across all triangles enables the evaluation of network stability and instability. Given the negative sign in the equation, a network with a predominancy of balanced (stable) triangles and well-coordinated organization of connections manifests as having negative energy. Conversely, a network with a predominancy of imbalanced (unstable) triangles and conflicting organization of connections is characterized by positive energy. This concept aligns with the principles of physical energy, where systems naturally tend toward greater stability at lower energy levels, while higher energy levels are generally associated with increased instability. In the equation, the combination term normalizes the energy between -1 and 1 by dividing by the number of triangles, where *N* is the number of regions, and (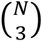) represents the number of triangles in a fully connected network. Including the cube root also facilitates a dimensionless representation, thus counteracting the redundancy introduced by multiplication.

Also, in the transition driven by a force, *F*, such as exposure to cognitive task, the network energy is altered as given by following equation, which is independent of the trajectory and intermediate states from rest to task assumed as a conservative process:

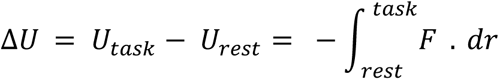

In previous works, we applied structural balance theory to the analysis of complex brain networks, conceptualizing it in the context of functional networks, studying patterns of structural balance, and exploring the significance of its application [10, 20-21]. We found that imbalanced triangles tend to form predominantly between canonical functional networks, while balanced triangles are more common within these networks during the resting state. Notably, subcortical regions play a significant role in the imbalance, whereas cortical areas, such as the visual cortex, are key in forming balanced triangles [20]. In another study, we highlighted the importance of structural balance by examining how the topology of negative functional connections, often overlooked, plays a critical role in the stability of the resting-state brain network. Our findings revealed that negative connections form hubs, pushing the network into a more stable state with lower energy than null networks with trivial topologies. This comparison validates the intrinsically efficient organization of brain connections during the resting-state [10]. In a lifespan study, we also discovered that imbalanced triangles, as the source of network instability and consequently network energy, follow a U-shaped pattern across lifespan in resting-state networks, reaching a minimum in early adulthood. This suggests increased brain network stability and well-coordinated network organization during early adulthood, when the brain has reached full maturity and is not engaged in development or degeneration [21].

Subsequently, numerous studies have explored the application of structural balance theory and network energy across various neural states and mental conditions. For instance, Talesh et al. observed a lower number of imbalanced triangles and a higher number of strongly balanced triangles in obsessive-compulsive disorder [22], leading to reduced network energy and potentially more stable brain states in this disorder compared to healthy controls. Similarly, Moradimanesh et al. reported reduced energy levels in both the salience network and the default mode network in autism spectrum disorder [23], which might be related to difficulties with dynamic switching and adaptability in behaviors, requiring further investigation. Fakhari et al. recently demonstrated a negative correlation between attention-deficit/hyperactivity disorder behavioral measures and brain network energy [24], suggesting that higher disorder severity may result in more stability in whole-brain resting-state network. Soleymani et al. found that perceiving pleasant stimuli is associated with lower energy and a more coordinated configuration, whereas unpleasant stimuli lead to higher energy and more conflicting network organization in the beta frequency band [25]. Additionally, Kashyap et al. used the concept of balanced and imbalanced triangles to evaluate the spread of stimulus effects following repetitive transcranial magnetic stimulation [26]. This suggests that network energy may serve as a valuable metric for assessing the efficacy of local brain stimulation. Overall, this line of research holds significant potential for advancing our understanding of neural mechanisms and therapeutic strategies. However, further efforts are needed to refine its framework, conceptualization, and interpretations, as well as to consolidate its validity and practical applications.

This study aimed to refine the conceptualization and interpretation of energy modeling in functional brain networks and extend its application to understanding the systematic organization of brain networks under cognitive control demands. We hypothesized that energy of whole-brain functional network would vary between resting state and tasks involving working memory, inhibitory control, and cognitive flexibility. Specifically, we sought to understand how energy is allocated across canonical functional networks depending on the nature of the task. Additionally, we investigated whether utilizing network energy as an informative input feature could improve predictive modeling performance. We also assessed the validity and reliability of the network energy measure to evaluate the robustness of this measure.

## Results

### Whole-brain network energy is altered in response to cognitive control tasks

The whole-brain functional network operates at different energy levels depending on cognitive control conditions. As shown in Figure 2, network energy significantly varies across task conditions (Friedman’s test for repeated measures, p < 0.001; non-parametric pairwise statistics are available in Supplementary Table 1). The task-free resting state, without specific cognitive demands, exhibits the lowest energy level, reflecting the optimal organization of functional connections across the brain and efficient system operating. Cognitive flexibility required during the shifting task brings the brain network to a lower energy level compared to other cognitive control conditions, suggesting a more coordinated whole-brain network organization in this task condition. Conversely, tasks involving inhibitory control (initiation and inhibition processes of go/no-go tasks, with fewer or more frequent go trials) and working memory (with different n-back orders) lead to higher energy levels in the whole-brain network. These task-dependent higher energy levels, compared to the task-free condition, highlight more conflicting functional connections’ organization and the enhanced adaptability in response to cognitive demands.

**Figure 2:**
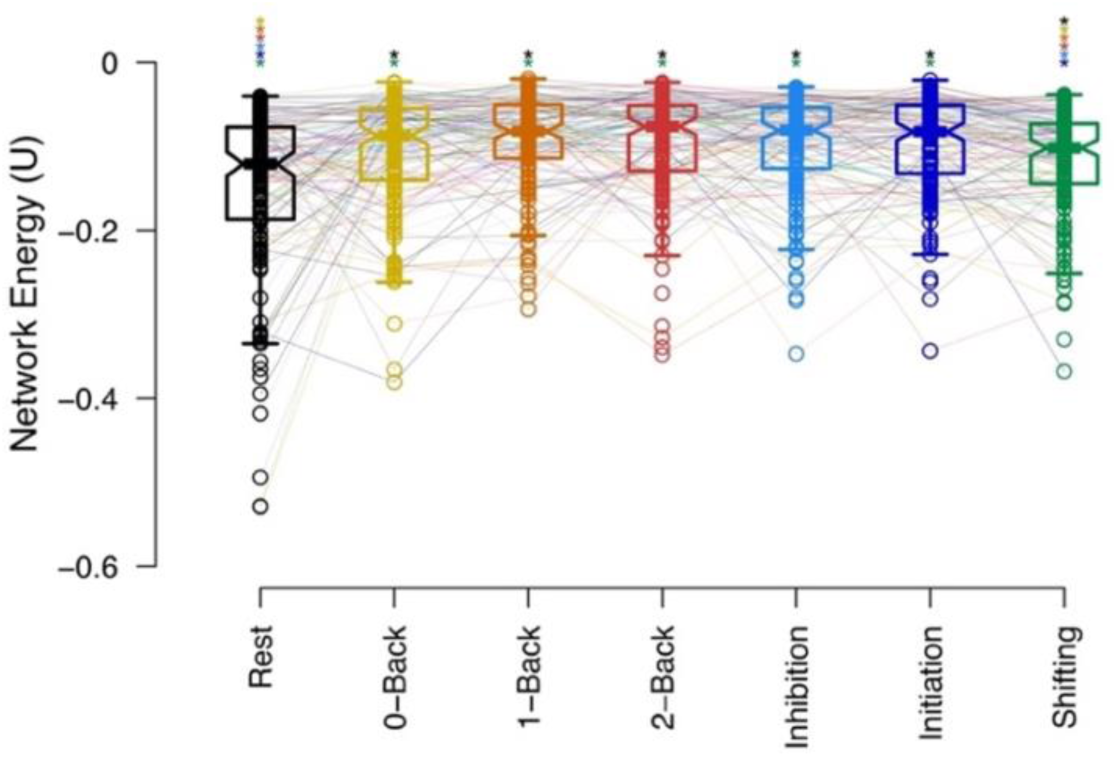
Whole-brain network energy during cognitive control tasks and resting state. Horizontal lines represent group-level medians, while notches on the boxes indicate the 95% confidence interval for the median. Individual subjects are linked by lines, with task conditions distinguished by color. Colored asterisks above the respective conditions denote significant corrected p-values for non-parametric pairwise comparisons between cognitive task conditions.

No significant differences were observed between energy levels for different difficulty levels of the n-back task or between the initiation and inhibition processes in the go/no-go task, suggesting that network energy reflects cognitive task types rather than the complexity level of a specific task.

Figure 2 presents whole-brain network energies derived using Power’s parcellation atlas [27], while Supplementary Figure 1 shows a replication of this analysis using Schaefer’s parcellation atlas [28].

### Canonical functional networks exhibit distinct patterns of energy distribution

Although the energy of the whole-brain network provides insights into the brain’s overall organization, examining energy within canonical functional networks, which are known for their involvement in specific sensory and cognitive functions, offers more precise insights into localized brain network organization. Figure 3A illustrates the diverse energy patterns across canonical functional networks during cognitive task conditions. The visual network consistently operates in lower energy levels, reflecting the coordinated organization of visually related regions, which are known for their effectiveness in processing sensory information. In contrast, the default mode network exhibits relatively elevated energy levels across all task conditions, highlighting its role in integrative brain and cognitive functions as a flexible, high-level network involved in various tasks.

**Figure 3:**
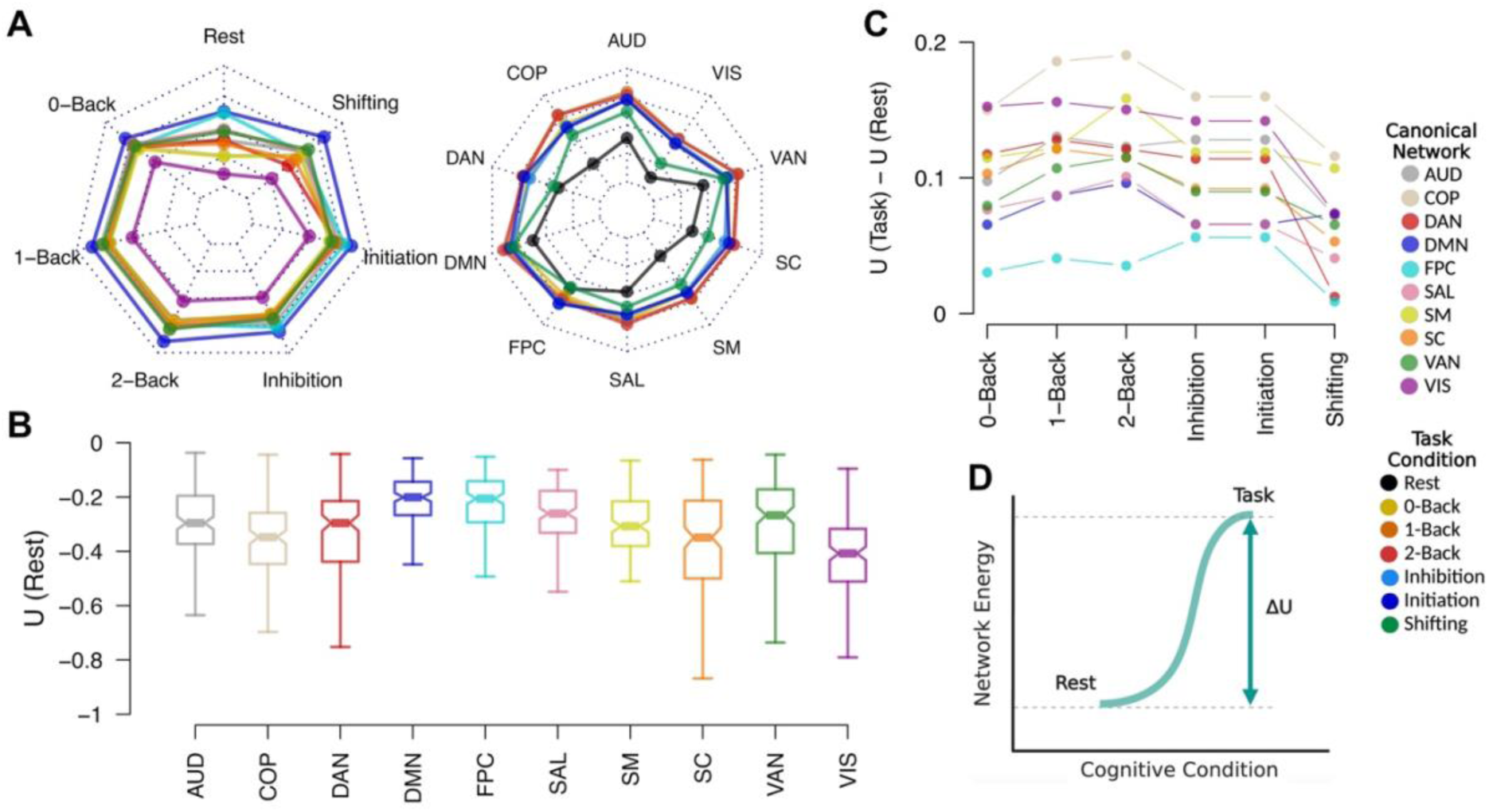
(A) Energy levels of canonical functional networks during resting state and cognitive control task conditions, shown as group-level medians. The color code on the right differentiates between cognitive task conditions and canonical networks. The radar chart displays energy values ranging between -0.45 and -0.05. (B) Energy levels of canonical networks during task-free resting state, with horizontal lines indicating group-level medians and notches representing the 95% confidence intervals for these medians. (C) Changes in network energy during the transition from rest to task state for each canonical network and cognitive task condition. Dots represent group-level medians of energy changes, with different networks indicated by colors. (D) A schematic representation of energy shifts during the transition from rest to task state. Abbreviations: AUD: auditory; COP: cingulo-opercular; DAN: dorsal attention; DMN: default mode; FPC: frontoparietal; SAL: salience; SM: somatomotor; SC: subcortical; VAN: ventral attention; VIS: visual; U: network energy; ΔU: network energy alteration.

Across all canonical networks, energy levels are lower during the resting state (Figure 3A right), indicating coordinated network organization in the absence of cognitive demands. Canonical networks associated with different cognitive task conditions exhibit distinct energy patterns. For instance, the dorsal attention network and subcortical structures show lower energy levels and more coordinated network organization during shifting tasks, highlighting their role in supporting cognitive flexibility.

Supplementary Tables 2–8 provide pairwise comparisons of canonical network energies across different task conditions, while Supplementary Tables 9–18 present differences in energy levels between task conditions for each individual network.

### Resting state shows greater network coordination in sensory systems and higher conflicting configurations in cognitive networks

Since our investigations indicate that the brain network operates at its lowest energy level during the resting state, with the most coordinated organization of functional connections, we focused on exploring this condition further. Figure 3B illustrates the energy levels of canonical networks during the resting state. The visual and subcortical networks, along with the somatosensory and auditory networks to a moderate extent, demonstrate lower energy levels. This reflects their efficiency in processing sensory input and likely supporting uniform homeostatic functions when the brain is at rest and free from external cognitive demands.

In contrast, the frontoparietal, default mode, salience, and ventral attention networks exhibit higher energy levels. These canonical networks are engaged in more complex and higher-order cognitive functions, including self-referential thought, attention regulation, and readiness to capture external stimuli. The elevated energy levels may represent the intrinsic flexibility required to transition between network configurations for broad processing of high-level cognitive functions. Although the cingulo-opercular network, which is mainly involved in cognitive processing, also shows lower energy levels, this reflects distinct mechanisms specific to this network.

Pairwise statistics comparing energy levels across canonical networks are presented in Supplementary Table 2.

### The energy levels of relevant canonical networks are purposefully changed during transitions from rest to cognitive task states

Although the energy of the whole-brain network increases during cognitive control tasks compared to rest, it is crucial to understand how the brain allocates this increased energy to the canonical functional networks. Therefore, we used the resting state, characterized by the most coordinated network organization and lowest energy, as a reference point and explored energy alterations in transition from rest to task across canonical networks (Figure 3.D). Selecting the resting state as a reference point aligns with its low energy level and its role in maintaining the subject’s readiness for exposure to cognitive tasks, as energy alterations reflect the demands imposed on the brain network during task engagement.

Figure 3.C illustrates the energy changes from the resting state to different task conditions. The visual and auditory networks, which exhibit the lowest energy levels during rest, experience the greatest energy increases when transitioning to task conditions. This increase reflects the need for these systems to become more flexible in order to receive and process sensory stimuli during task performance. In contrast, networks associated with higher-order cognitive processing (e.g., frontoparietal and default mode) receive significantly less energy, reflecting the efficient processing of these systems and less energy demand during cognitive tasks. This emphasizes the distinct energy allocations between sensory and cognitive systems during the transition from rest to task.

In the n-back task, the frontoparietal network received only a minor increase in energy compared to other networks. This suggests that this system operates more efficiently to support working memory demands than other networks. Consequently, it does not require usual share of energy, and the increased energy associated with the transition from rest to task is distributed across other networks. Supplementary Tables 19–21 provide non-parametric statistical comparisons of energy changes between canonical networks across various n-back conditions.

In the transition from the resting state to the inhibitory control task, the energy increase across canonical networks appears more uniform, with no significant differences observed between the initiation and inhibition conditions, despite the unequal proportion of go/no-go trials. Specifically, the frontoparietal, default mode, and salience networks, crucial for the cognitive processing required for inhibitory control, receive less energy, emphasizing their efficiency during the go/no-go task. Supplementary Tables 22–23 provide non-parametric statistical comparisons between canonical networks across the initiation and inhibition conditions of the go/no-go task.

Figure 3.C also shows that both the frontoparietal and dorsal attention networks, important for cognitive flexibility, receive smaller amounts of whole-brain elevated energy in transition from rest to shifting task, reflecting their efficient operation in supporting this cognitive function. A moderate version of this effect is also observed in the salience network and subcortical structures. Statistical details for pairwise comparisons among canonical networks can be found in Supplementary Table 24.

### Network energy, as an input feature, enhances the performance of cognitive control task classification and predicts chronological age

In the previous sections, we demonstrated the application of network energy in exploring network organization, which reflects the flexibility and efficiency of brain systems across cognitive conditions and during transitions from rest to task. To further investigate the potential of network energy as a global network measure, we designed a series of predictive models aimed at classifying cognitive control task states and predicting chronological age. Our objective was to evaluate the effectiveness of brain network energy as an input feature for predictive modeling, compared to other established global network metrics, such as the global clustering coefficient, global efficiency, and modularity. The analysis included network measures extracted from both the whole brain and canonical networks as inputs for the models. We employed a subject-wise leave-one-out cross-validation approach with support vector machine using radial kernel for predictive modeling.

For the classification modeling, we aimed to discriminate between four cognitive control task states: working memory, cognitive flexibility, inhibitory control, and task-free resting state. Figure 4 presents the results of cognitive task classification using different feature types as model inputs. Figure 4A shows the classifier’s accuracy for the overall model and each cognitive task state, based on modeling with various global network measure types derived from whole-brain and canonical networks, as well as combinations of all network measure types.

**Figure 4:**
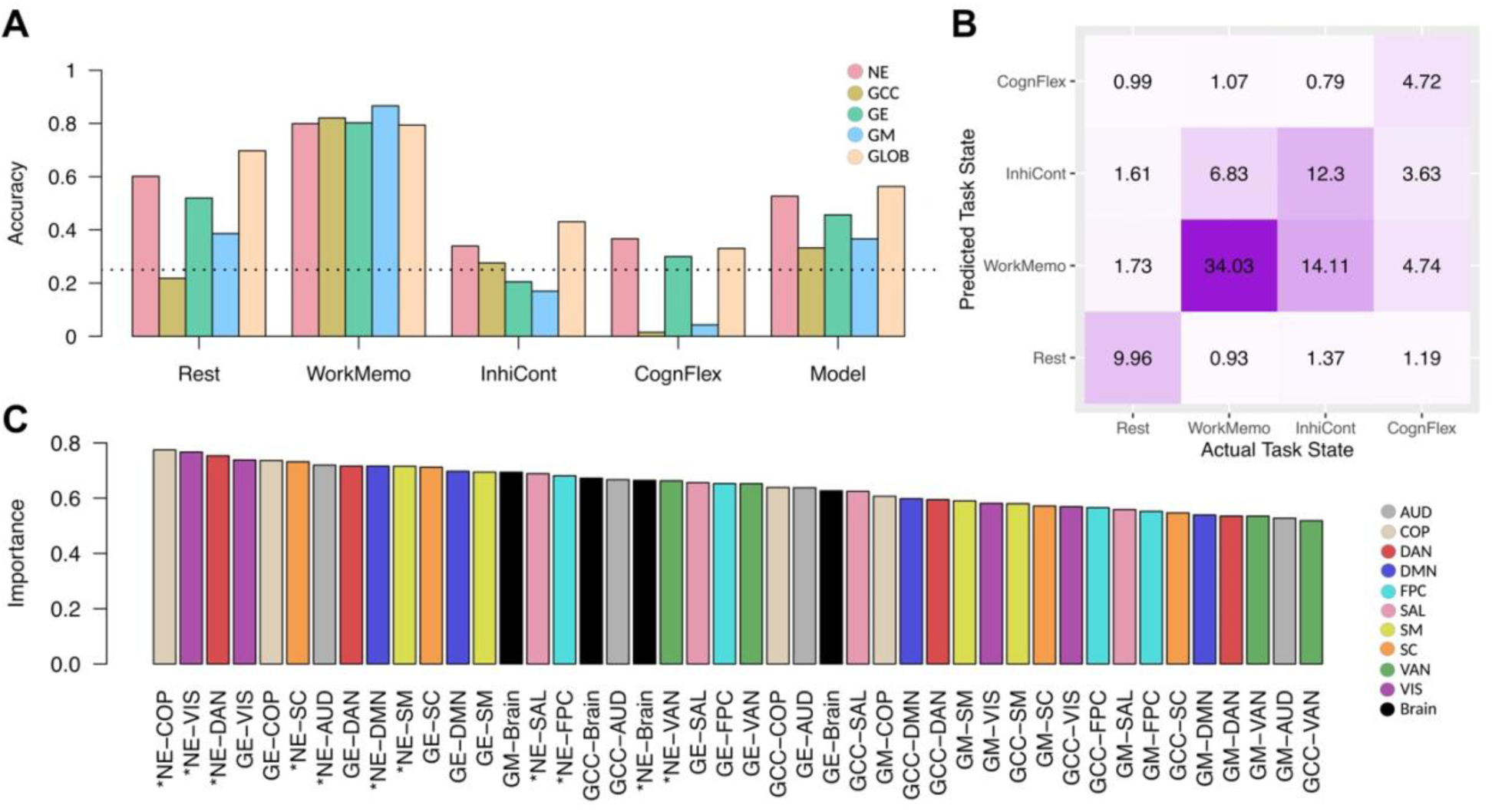
Classification of cognitive control task states. (A) Bar plots display the classifier’s accuracy for each cognitive task state (class) individually, along with the balanced accuracy for the overall classification model. The colors of the bars represent different subsets of input features. The horizontal dotted line marks the chance level of classification. (B) Confusion matrix for the classification model using all network measure types as input features during training. The numbers indicate the percentage of actual values that were predicted for each class, with darker cell colors representing higher values. (C) Bar plot illustrates the importance of different features in cognitive task state classification. The colors of the bars represent various canonical networks. Asterisks on the x-axis labels denote network energy features extracted from functional networks. Abbreviations: NE: network energy; GCC: global clustering coefficient; GE: global efficiency; GM: global modularity; GLOB: all global network measure types; AUD: auditory; COP: cingulo-opercular; DAN: dorsal attention; DMN: default mode; FPC: frontoparietal; SAL: salience; SM: somatomotor; SC: subcortical; VAN: ventral attention; VIS: visual.

Notably, using network energy as an input feature improved the discrimination of cognitive task states (balanced accuracy = 0.53), achieving the highest per-state accuracy, except for the working memory task, compared to other global network measure types. As expected, the classifier achieved the highest overall accuracy when incorporating all global network measure types as input features (balanced accuracy = 0.56). Detailed accuracy values across cognitive task classes and network measure types are provided in Supplementary Table 25. Figure 4B presents the confusion matrix for the cognitive task state classification model trained with all global network measure types. Finally, Figure 4C highlights the importance of various features in the classification model, showing that network energies from both canonical and whole-brain networks are the most influential, as indicated by asterisks on the x-axis.

In our regression modeling, we aimed to predict chronological age based on various global network measure types derived from whole-brain and canonical networks across both resting-state and cognitive tasks. To evaluate the predictive capacity of network energy compared to other global network measure types, we used them as model inputs (Figure 5). Specifically, Figure 5A shows that the model performed better and reduced prediction error when network energy was used as the input feature (R² = 0.447, MAE = 11.29) compared to global clustering, efficiency, and modularity metrics. The best age prediction performance was achieved when all global network measure types were combined as inputs (R² = 0.547, MAE = 10.36). Supplementary Table 26 provides detailed performance metrics for the regressor using each global network measure type and their combinations. Figure 5B presents the correlation between predicted and actual ages for the model incorporating all graph measure types. Figure 5C further highlights the importance of network energy features in the model’s performance, with their significance indicated by asterisks on the x-axis. Notably, this figure also emphasizes that network measures from the cingulo-opercular network were particularly crucial for age prediction. Overall, these predictive models underscore the significance of network energy as a key global network measure in machine learning models applied to brain and cognitive research.

**Figure 5:**
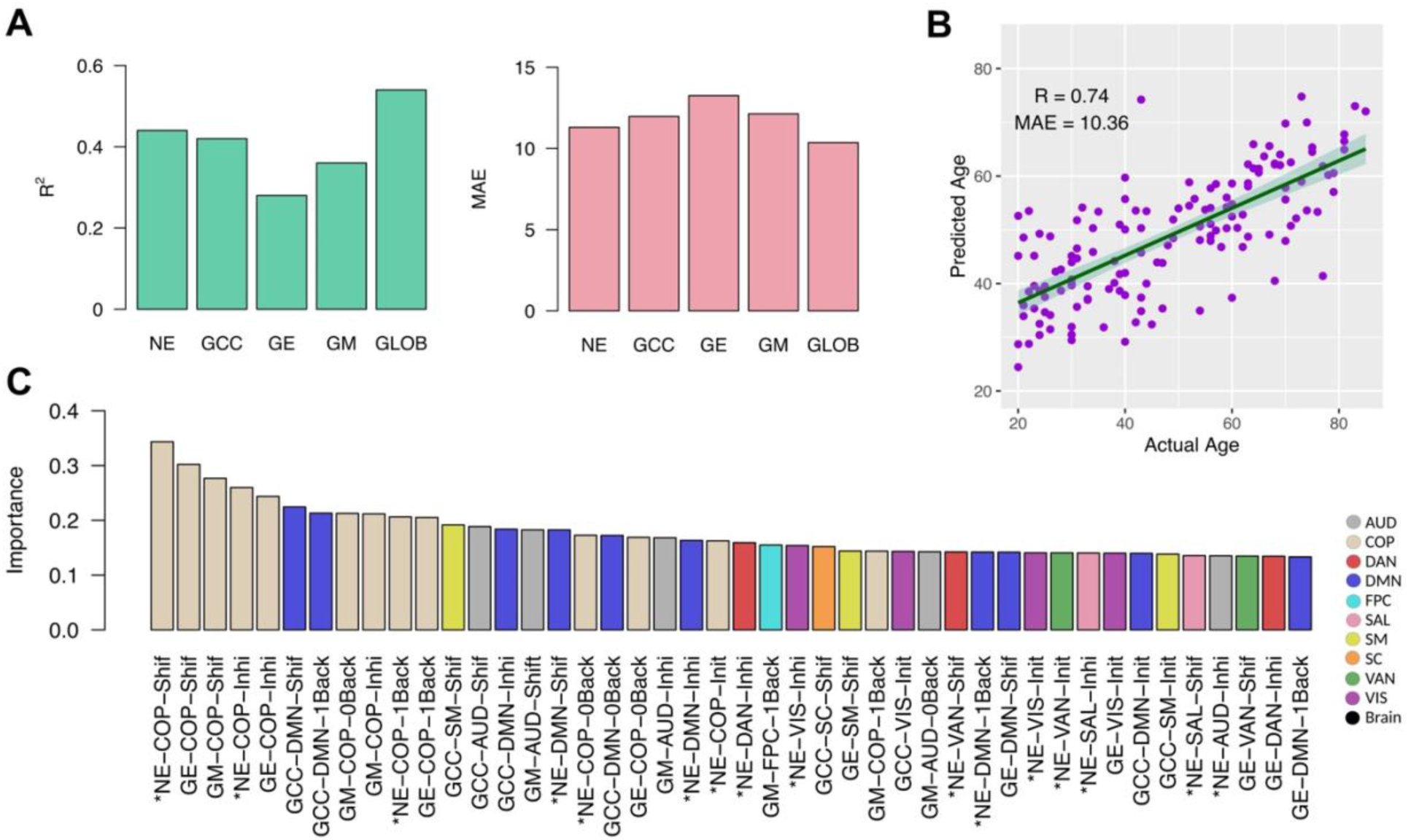
Regression modeling of chronological age. (A) Bar plots show the performance and error in age prediction for various subsets of network measures utilized as input features. (B) Scatter plot demonstrates actual versus predicted age using the full set of input features obtained from all network measures, with each dot representing an individual subject and the fitted line showing the linear relationship. (C) Bar plot illustrates the importance of different features in predicting chronological age when we trained the model using all network measure types. The colors of the bars represent various canonical networks, with features related to network energy marked by asterisks on the x-axis labels. Abbreviations: R²: coefficient of determination; MAE: mean absolute error; NE: network energy; GCC: global clustering coefficient; GE: global efficiency; GM: global modularity; GLOB: all global network measure types; AUD: auditory; COP: cingulo-opercular; DAN: dorsal attention; DMN: default mode; FPC: frontoparietal; SAL: salience; SM: somatomotor; SC: subcortical; VAN: ventral attention; VIS: visual.

### Network energy serves as a valid and reliable measure

To investigate the validity of network energy measurements and ensure that our results are not a consequence of random network organization, we compared whole-brain network energy during various cognitive tasks and the resting state with corresponding null networks that have random topologies. As illustrated in Figure 6A, the actual brain networks during cognitive control tasks and resting state exhibit significantly lower energy levels compared to the null networks. These findings suggest an intrinsically more coordinated (less conflicting) organization of functional connections and reduced network energy, underscoring the non-random nature of the observed network energies.

**Figure 6:**
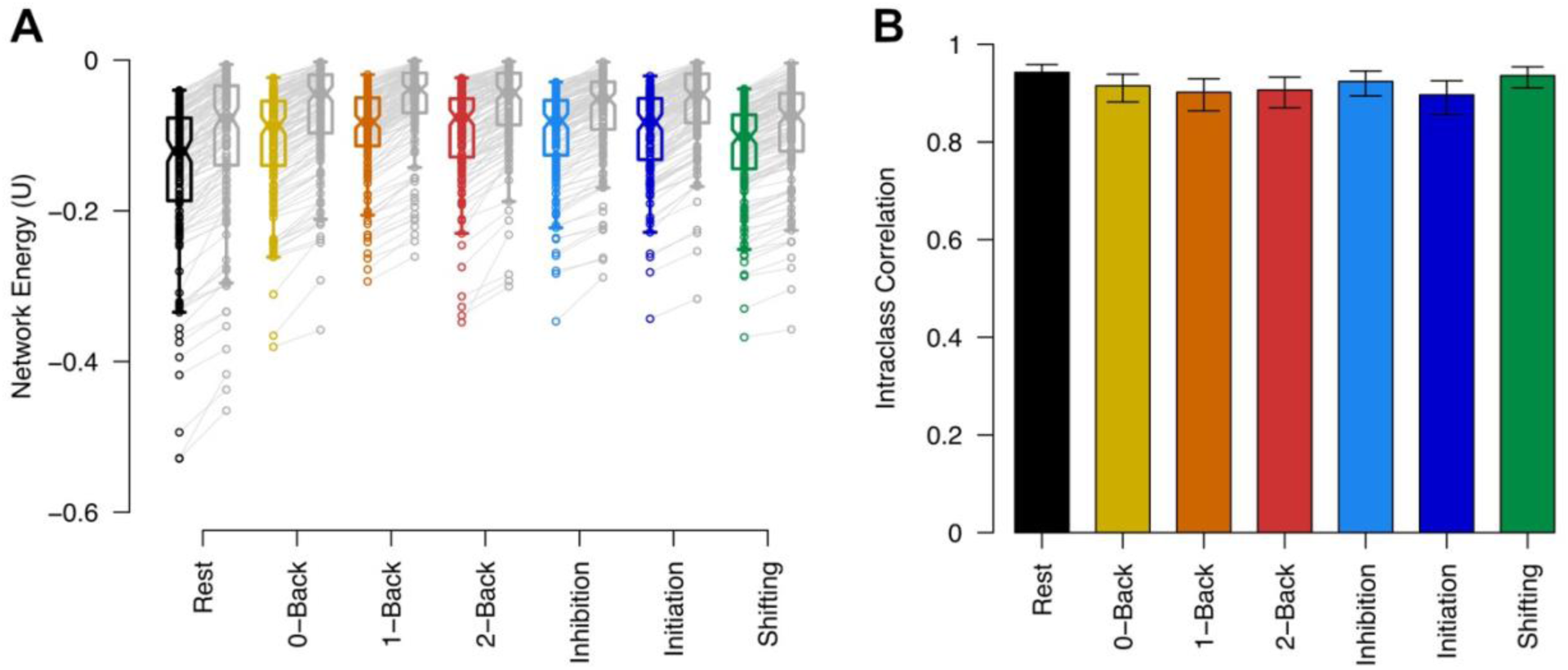
Validity and reliability of network energy. (A) Energy comparison between actual whole-brain networks and their corresponding null networks with random topologies. Horizontal lines in the box plots denote group-level medians of energy levels, and notches represent the 95% confidence intervals. Actual networks corresponding to cognitive task conditions are color-coded, while the related null networks are shown in gray, adjacent to each task condition. Lines connect each subject’s network to its corresponding null network. (B) Intraclass correlation of whole-brain network energies between networks formed using Power’s atlas and Schaefer’s parcellation atlas for each cognitive task condition. The height of the bar plots indicates the average fixed raters’ ICC values, with error bars representing the corresponding 95% confidence intervals. Colors distinguish between task conditions.

Additionally, the brain network energy is not at the absolute stable state with the lowest energy level, supporting the idea that the brain minimizes energy for optimized processing while maintaining flexibility. Supplementary Table 27 provides the corresponding p-values and statistics related to the Wilcoxon signed-rank test between the energy of actual and null networks for each cognitive task condition.

To evaluate the reliability of network energy measurements, we computed the energy of the whole-brain network using Schaefer’s atlas for each cognitive task condition and compared the results with network energies calculated using Power’s parcellation atlas (Figure 6B). The intraclass correlation analysis indicated a high degree of reliability between the results obtained from the two atlases for each cognitive task condition. This strong correlation suggests that the network energy measure is not dependent on the specific brain parcellation used. The average intraclass correlation values for fixed raters, along with the corresponding confidence intervals, are presented in Supplementary Table 28.

## Discussion

### Summary

Our study provided valuable insights into how network energy is distributed across functional brain networks during cognitive control tasks, demonstrating coordinated or conflicting configurations of functional connections that reflect the efficiency and flexibility of brain systems.

The resting state, characterized by the lowest energy level, indicates a highly stable and coordinated whole-brain organization of functional connections, whereas cognitive control tasks demonstrate increased energy levels, reflecting greater instability and conflicting network organization (Figure 2). These findings suggest that cognitive demands are associated with more adaptability in the whole-brain system, while the task-free state demonstrates more proportionally efficient processing. No significant differences were observed in energy levels across the different difficulty levels of the n-back task or between initiation and inhibition conditions in the go/no-go task, indicating that network energy primarily reflects task type differences rather than the complexity of a specific task. Sensory networks, such as the visual and somatosensory networks, along with the subcortical network, exhibited relatively lower energy levels during the resting state, demonstrating their coordinated organization and reflecting more efficient processing in the absence of external stimuli. In contrast, networks involved in cognitive functioning, such as the default mode network, frontoparietal network, and salience network, exhibited relatively higher energy levels and more conflicting network organization during the resting state. This may correspond to their intrinsic flexibility, enabling them to support broad high-level cognitive processing (Figure 3B).

Although involvement in cognitive control tasks increases whole-brain energy (Figure 2), this increased energy is not allocated uniformly across canonical functional networks (Figure 3A). During the transition from rest to task conditions, the visual and auditory networks receive relatively more energy, indicating greater conflict in network configurations, which reflects increased flexibility to process broad incoming sensory stimuli (Figure 3C). In contrast, cognitive networks, such as the frontoparietal and default mode networks, receive only a minor portion of the increased whole-brain energy, suggesting their optimized functioning and less energy demands under cognitive control. This efficiency is particularly evident in canonical networks known for processing specific cognitive functions, such as the frontoparietal network in working memory tasks. These findings emphasize distinct mechanisms of energy allocation between sensory and cognitive systems during cognitive control.

Our predictive modeling demonstrated the utility of network energy as an input feature type in machine learning approaches (Figure 4 and Figure 5). Utilizing network energy features extracted from functionals network across cognitive conditions significantly improved the performance of the classification of cognitive task states and prediction of chronological age. These results suggest that network energy not only reflects the efficiency and flexibility of brain systems but also serves as an informative global network measure for developing biomarkers in the brain studies.

Lastly, our validation and reliability analyses confirmed that the observed network energy measurements are not random effects (Figure 6A). The lower energy levels of actual brain networks compared to null networks with random topologies underscored the brain’s intrinsic coordination of functional connections. Additionally, the strong agreement between energy measurements across different brain parcellation schemes, Power’s and Schaefer’s atlases, supported the robustness of this metric, with intraclass correlation analysis indicating high reliability.

### Structural balance theory in brain networks: A valid analogy?

One important consideration in applying the structural balance framework is the analogy between social networks and brain connectivity. In social contexts, interpersonal relationships, attitude changes, persuasion, and social influence are significantly influenced by higher-order interactions among interconnected entities. This inherent interdependence is precisely what Fritz Heider considered when theorizing balance theory [11-12].

Similarly, the interdependence of brain regions and functional connections strengthens the validity of applying structural balance theory to brain networks. Shared information and dependencies between brain components could give rise to non-trivial organization within brain networks, as we observed in the comparison of actual and null networks (Figure 6A).

### Resting state beyond a single minimal energy configuration

Our analysis suggests that the brain operates at relatively low energy levels during rest, consistent with previous research demonstrating that a broad range of resting-state patterns converge toward low-energy configurations [3]. However, it is essential to recognize that the resting state does not represent a singular ‘minimal energy state.’ Instead, behavioral and neuroimaging studies support the existence of multiple low-energy states, encompassing a spectrum of transient brain states [2, 4-5]. Thus, our findings should not be interpreted as evidence of a single minimal energy configuration but rather as a reflection of the brain’s well-coordinated network organization in the absence of external task demands. Our results may present the resting state as a metastable state, ready to transition to higher energy states involving cognitive control. This underscores the crucial role of the resting state as a central reference point in brain energy studies.

### Challenges in mapping the energy landscape of brain networks

While our study focuses on exploring network energy across cognitive tasks and resting state, it is not a detailed analysis of the energy landscape. Specifically, we did not directly analyze the presence of attractors, saddle points, or energy barriers, which are essential components of an energy landscape framework. These elements are crucial for precisely identifying the stability and transitions between brain states, as they provide deeper insights into how the brain shifts between various cognitive states and maintains equilibrium within multi-stable regimes. To explore the brain network’s energy landscape more thoroughly, methodologies from previous works, such as those by Watanabe et al. (2013), Ashourvan et al. (2017), and subsequent studies, can be adopted [1-9, 29]. These works analyzed energy landscapes in terms of stable states (attractors), transitions between states, and the energy barriers that separate them.

Furthermore, while our findings describe network energy differences between rest and task conditions, they do not provide detailed information on the transition pathways, intermediate states, or energy barriers involved in these shifts. We interpreted these changes as transitions because, in physics, it is possible to gain general insights into transitions by comparing the energy of the initial and final states without considering the specific path taken between them in a conservative process.

It is obvious that our study is just a starting point. Future studies could address this limitation by utilizing dynamic functional connectivity [30] or hidden Markov models [31] to track the brain network’s intermediate states as it transitions between rest and task conditions. This approach would provide a more continuous understanding of state changes, as well as the energy barriers that need to be overcome for the brain to shift from one cognitive state to another. By analyzing the stability of different brain network states and the transitions between them, we could offer a more robust interpretation of the energy landscape and how it governs brain network dynamics during cognitive tasks.

### Energy definition: a comparison with the maximum entropy model

The energy function conceptualized and introduced by Watanabe et al. (2013), based on the maximum entropy model, provides a valuable framework for exploring brain states, basins, and state transitions through the statistical evaluation of observed states [1]. This model has been widely applied in brain energy landscape studies and offers significant insights [2-9]. However, there are some limitations to consider.

While this model utilizes the time course of functional images to estimate energy parameters, it defines states and energies for each brief scanning period. Given the inherent low signal-to-noise ratio of fMRI data, this method may raise concerns about the validity of the resulting brain energy estimates. Additionally, the model binarizes the activation of brain regions to estimate energy, potentially losing information about activation dynamics.

In contrast, our approach does not rely on instantaneous activations but instead uses the full time-course of brain activations and their coactivations to assess energy over a longer recording period. This avoids potential issues related to the low signal-to-noise ratio of fMRI signals. Metaphorically, our method evaluates the average energy over a cognitive condition period, while Watanabe et al.’s method computes instantaneous energies. Although our approach limits the ability to investigate transition paths between states, this limitation could be addressed in future work by employing dynamic brain networks with carefully selected sliding window widths.

Moreover, our approach does not binarize activations, instead considering weighted connections to preserve all available information in energy calculations. Our analysis demonstrated the reliability and validity of the calculated energies (Figure 6).

It is also important to note that our energy function specifically addresses the organization of functional brain networks, in contrast to Watanabe’s model, which explores the energy of the entire brain system. The network energy metric in this study also does not refer to the metabolic processes of the brain.

### Balanced/imbalanced triangles versus network motifs: complementary insights into brain network functionality

Network energy is derived based on the accumulation of balanced and imbalanced triangles formed by three connected brain regions, providing insights into the efficiency of network organization. However, in neuroscience, the concept of network motifs is another widely discussed feature for understanding the structure and function of brain networks.

Network motifs are defined as small, recurring patterns of interconnections that occur more frequently than expected by chance in a given network. These motifs serve as fundamental building blocks for complex networks and have been extensively studied in brain network research [32-34]. Motifs such as feedforward and feedback loops are crucial for understanding information flow in brain networks.

While both network energy and motifs refer to small-scale structures within networks, they capture different aspects of brain network functionality. Balanced and imbalanced triangles reflect the stability and instability of relationships, particularly with respect to the positive or negative signs of connections, which represent coordinated or conflicting higher-order interactions. Motifs, on the other hand, do not necessarily account for the signs of connections but rather focus on specific structural patterns of binary connections that are dominant in brain networks. Although a motif may appear structurally similar to a triangle analyzed in our energy framework, but the focus differs from stability and flexibility to the role that configuration plays in pattern frequency and information processing.

### Alignment of energy allocations with established roles of cognitive networks

In the transition from the resting state to cognitive task conditions, accompanied by elevated whole-brain energy, our findings highlight distinct energy alterations between cognitive and sensory networks. Cognitive networks, well-known for their involvement in cognitive functioning, require only minimal energy elevation. In contrast, sensory networks receive relatively higher energy levels. We interpret this as a reflection of the fact that key cognitive systems, operating efficiently, do not require a equal share of this elevated energy. Conversely, sensory systems, which interact directly with the environment, need to maintain flexibility, reflecting induced conflicting configurations, and therefore receive a larger portion of the elevated energy.

Our results reveal that the frontoparietal network exhibits only a minor energy increase from rest to the n-back task. This finding aligns with prior studies that emphasize the frontoparietal network’s role in supporting working memory and cognitive control processes [35–40]. The minimal network energy increase suggests that the frontoparietal network operates with optimized functionality, effectively managing working memory demands without requiring significant energy input compared to other networks.

In the transition from the resting state to the inhibitory control task, the frontoparietal, default mode, and salience networks exhibit smaller energy increases, with the ventral attention and subcortical networks showing a slightly weaker version of this effect. The efficiency of processing in these systems, inferred from their minimal energy increases relative to the rest of the brain, aligns with the active role of frontostriatal pathways and subcortical structures in supporting action inhibition, as well as the importance of default mode network reorganization for successful inhibitory control [41–51]. This interpretation highlights the efficient functionality of these network systems during working memory task, enabling them to meet task demands without requiring a equal share of the elevated whole-brain energy.

From rest to the cognitive flexibility condition, the frontoparietal network and the dorsal attention network, which mediate top-down attention, exhibit the smallest energy increases. Other networks, such as the default mode, subcortical, and bottom-up attentional networks, show moderate energy increases. The efficiency of processing in these systems during cognitive flexibility, inferred from their minimal allocation of whole-brain energy, aligns with their involvement in cognitive adaptability, attention shifts, and behavioral modulations [52–60]. These findings underline the capacity of these network systems to fulfill cognitive demands effectively, with only minor utilization of the elevated whole-brain energy.

### Distinct role of the cingulo-opercular network in cognitive control and aging

Our study identified a distinct pattern of energy allocation in the cingulo-opercular network during cognitive control tasks (Figure 3.C), which contrasts with other cognitive networks that exhibit only minor energy increases due to their efficient processing. This suggests a unique role for the cingulo-opercular network in cognitive control, as previous research has consistently demonstrated its involvement in working memory, concurrent cognitive control, and task switching [61–63]. Further investigation is needed to better understand the mechanisms underlying the systematic organization of this canonical network in cognitive control.

Additionally, our results highlight the relevance of the cingulo-opercular network in aging, with network features obtained from this network being crucial for age prediction through regression models (Figure 5.C). This aligns with previous research indicating age-related declines in functional connectivity within the cingulo-opercular network. Notably, studies have linked these declines to a range of cognitive impairments, including global cognitive states, visuospatial abilities, and executive functions [64–68]. Collectively, these findings underscore the critical role of the cingulo-opercular network in cognitive aging and its potential in developing biomarkers for age-related cognitive decline.

### Improving predictive models by employing network energy features

Network measures, combined with machine learning, have played a significant role in understanding neurological processes and disorders [69]. Predictive models utilizing network measures as input features have been successfully applied in clinical research, including studies on Parkinson’s disease, autism spectrum disorder, and Alzheimer’s disease, yielding promising diagnostic results [70-72]. This approach has also been used to distinguish emotional states, analyze neural network changes in epilepsy, and predict treatment outcomes in schizophrenia [73-75].

Our study presents network energy as an informative global network feature for machine learning applications in neuroscience (Figures 4 and Figure 5). Our findings suggest that network energy, as a measure of both the coordinated and conflicting network organization, can enhance model performance, including predicting age and distinguishing between cognitive states. This offers promising broader applicability for network energy in various neurological conditions.

### Limitations and future directions

Our study offers valuable insights into the organization of functional connections through network energy across various cognitive task conditions, but it has some limitations. First, our analysis did not incorporate a detailed exploration of the brain’s energy landscape, such as identifying attractors, saddle points, or energy barriers, which are critical components for understanding transitions between brain states. Without this level of detail, our understanding of how the brain shifts between cognitive states remains incomplete. Future studies should adopt more refined energy landscape frameworks.

Additionally, our approach does not directly track dynamic changes over time within network configurations, limiting our ability to explore transition pathways or intermediate states between rest and task conditions. Implementing dynamic functional connectivity models or hidden Markov models in future research could address this limitation by providing a more continuous view of the brain network’s energy shifts during different cognitive tasks.

Although we validated network energy as a reliable measure using two brain parcellation schemes (Power and Schaefer atlases), further replication with other parcellation approaches and across more diverse datasets is necessary to ensure the generalizability of our findings.

Future research could also integrate more individual-specific analyses, particularly in relation to aging, cognitive performance, and neuropsychiatric disorders. Personalized network energy metrics could help predict clinical measures, offering valuable biomarkers for clinical research. Expanding the application of network energy as a machine learning feature to domains such as neurological diseases, neurodevelopmental disorders, and mental health conditions could enhance the performance of diagnostic tools. Future studies could also explore how network energy patterns differ in response to neural interventions, broadening the clinical application of this measure.

## Conclusion

In conclusion, our study highlights the selective allocation of energy to functional brain networks across different cognitive tasks, demonstrating distinct patterns of energy distribution in response to cognitive demands. The whole-brain exhibits more coordinated network organization with lower energy level during task-free condition, whereas cognitive control tasks are associated with more conflicting network organization and elevated energy levels. Notably, sensory networks receive the largest share of the elevated whole-brain energy when transitioning from rest to task conditions, reflecting their flexibility in adapting to and processing external stimuli. In contrast, cognitive networks operate more efficiently, requiring a smaller proportion of this elevated energy.

Our findings also underscore the potential of network energy as a significant global network measure in predictive models. Furthermore, the validity of energy measurements obtained from comparisons to null models, along with the strong reliability of network energy measurements across different brain parcellation schemes, supports the robustness of network energy and highlights its utility in developing biomarkers.

## Methods

### Dataset

In this study, we utilized a publicly available functional connectivity dataset provided by J.R. Rieck et al. [76]. The dataset comprises 144 healthy participants, aged 20–86, from the Greater Toronto Area. These participants underwent fMRI scans while engaging in three distinct cognitive control tasks, including working memory, inhibitory control, and cognitive flexibility tasks, as well as a resting-state scan. The data were acquired using a Siemens Trio 3T magnet at Baycrest Health Sciences. Informed consent was obtained from the participants in accordance with a protocol approved by the Research Ethics Board at Baycrest Health Sciences Center.

Following standard preprocessing and postprocessing, the dataset offers seven connectivity matrices for each subject: three corresponding to the 0-, 1-, and 2-back tasks of working memory; two for the inhibition and initiation conditions of the go/no-go task, with corresponding most and less go trials for inhibitory control; one for the shifting task of cognitive flexibility; and one for the task-free resting-state condition. The connectivity profiles were derived based on Power’s and Schaefer’s atlases [27-28]. We utilized Power’s parcellation atlas, which divides the brain into 229 regions across ten canonical functional networks, in the main analysis and Schaefer’s atlas with 200 cortical regions for the evaluation of the reliability of results. So we had a 229 × 229 whole-brain connectivity matrix (or a 200 × 200 Schaefer’s matrix) for each cognitive task condition of each subject, containing pairwise correlations between regional activations of each pair of regions. We utilized subsections of this whole-brain matrix for each canonical network, with the ROIs belonging to their respective canonical networks. The coordinates of the ROIs and their corresponding canonical network labels are available in the dataset.

For further information regarding the procedures of the cognitive tasks and the employed fMRI data processing, you can refer to the original reference paper by J.R. Rieck et al. [76].

### Statistical analysis

We calculated the network energy of the whole-brain network and individual canonical networks for each cognitive task condition of each subject. Due to the non-normal group-level distribution of network energy measures, both at the whole-brain and canonical network levels, we employed non-parametric statistical methods for our analysis. The Friedman repeated measures test was used to compare network energy across various cognitive task conditions (Figures 2 and Figure 3). Additionally, the Wilcoxon signed-rank test was employed for pairwise post-hoc comparisons, with multiple comparison corrections applied (Figures 2 and Figure 3). We also used the Wilcoxon signed-rank test to compare the energy values of actual networks with energy of corresponding null networks with random topology for each cognitive task condition (Figure 6A). For the reliability analysis of network energies derived from Power’s and Schaefer’s atlases, we utilized intraclass correlation analysis (Figure 6B).

### Predictive modeling

We employed the support vector machine with radial basis function kernel for two main purposes: classifying different cognitive task states and predicting age (Figures 4 and 5). To ensure accurate and reliable model performance, we used a participant-wise leave-one-out cross-validation technique, which effectively prevents overfitting. During model training, we performed resampling over various tuning parameters, holding the ‘sigma’ parameter constant while fine-tuning the regularization parameter ‘C’. The optimal model was selected based on the highest accuracy for classification modeling and the lowest mean absolute error for regression modeling.

We computed three global network measures: global clustering coefficient, global efficiency, and global modularity, as well as network energy for the whole-brain and canonical networks for each cognitive conditions of each subject. We then explored the predictive power of these measures by using subsets of each measure type derived from whole-brain and canonical networks during cognitive tasks and resting-state conditions as inputs for our predictive models. The performance of models based on different network measure types was compared to assess the predictive effectiveness of each network measure type. To further enhance model interpretability, we performed predictive modeling based on all network measure types across all functional networks, which improved performance. Then we evaluated the significance of each network measure corresponding to a brain network and cognitive condition by analyzing feature importance based on the magnitude of standardized coefficients.

### Computational and graphical tools

All computations, including the extraction of network measures, null modeling, statistical comparisons, machine learning modeling, and figure generation, were performed using R and its associated packages [77-83].

## Data availability

The original dataset is accessed at https://osf.io/m5crs/, and our codes are hosted on GitHub at https://github.com/majidsaberi/NetEnergyCogControl for anyone interested in implementing, replicating, or further developing our work.

## Glossary

Structural balance: A theoretical framework used to assess the stability or instability of network organization by categorizing triangles into balanced or imbalanced configurations. These classifications are then used to calculate the network’s energy, which reflects the overall state of coordination or conflict within the network.
Network energy: A metric derived from the structural balance framework that quantifies the level of conflict or coordination in the configuration of connections within a network. It reflects the network’s overall stability or instability by measuring the proportion and intensity of balanced and imbalanced triangles.
Balanced/imbalanced triangle: These terms describe the arrangement of connections within a triangle in a network. A balanced triangle is stable, with relationships coordinated to minimize tension among components, promoting overall network stability. In contrast, an imbalanced triangle is unstable, representing conflict among components and requiring reorganization to achieve stability. Originally defined in the context of personal relationships, a triangle is considered balanced when three or one of the relationships are friendships (e.g., a "friend of my friend" or an "enemy of my enemy" is a friend). Conversely, a triangle is imbalanced when three or one of the relationships are hostile (e.g., a "friend of my friend" or an "enemy of my enemy" is an enemy), generating tension within the system.
Conflicting network organization: A network configuration characterized by high levels of tension or dissonance among connections. It is associated with an increased proportion and intensity of imbalanced triangles, and elevated network energy levels, reflecting instability within the network.
Coordinated network organization: A network configuration characterized by an optimal arrangement of connections that minimizes conflict, resulting in stability and coordination. It is associated with a high proportion and intensity of balanced triangles and low levels of network energy.
System flexibility: The ability of the brain system to adjust and adapt in response to external demands. It is associated with increased network energy requirements and dynamic reorganization of connections.
System efficiency: It refers to the optimal functionality of a brain system specialized for specific neural processing. It is associated with reduced network energy requirements for effective operation.
Energy allocation: The process by which the brain distributes energy across different canonical networks, depending on task demands, the specialization of the networks, and their corresponding requirements for flexibility or efficient processing.
Canonical functional networks: Predefined, large-scale networks in the brain (e.g., default mode, dorsal attention, visual networks) associated with specific sensory or cognitive processes.

## Author contributions

The study was conceptualized by MS and AK. Supervision was provided by AK, BTD, and BM. Data analysis was performed by MS, while original data collection and processing were carried out by JRR and CLG. MS was responsible for the visualization and creation of figures. The original draft of the manuscript was written by MS, and all authors contributed to reviewing and editing the manuscript.

## Acknowledgment

First author would like to express his gratitude to Maryam Azimi for her insightful feedback on the manuscript and her invaluable support throughout the research process.

## References

1. Watanabe T, Hirose S, Wada H, Imai Y, Machida T, Shirouzu I, Konishi S, Miyashita Y, Masuda N. A pairwise maximum entropy model accurately describes resting-state human brain networks. Nature communications. 2013 Jan 22;4(1):1370.

2. Gu, S. et al. The Energy Landscape of Neurophysiological Activity Implicit in Brain Network Structure. Scientific Reports vol. 8 (2018).

3. Watanabe, T. et al. Energy landscapes of resting-state brain networks. Frontiers in Neuroinformatics vol. 8 (2014).

4. Watanabe, T., Masuda, N., Megumi, F., Kanai, R. & Rees, G. Energy landscape and dynamics of brain activity during human bistable perception. Nature Communications vol. 5 (2014).

5. Ashourvan, A., Gu, S., Mattar, M. G., Vettel, J. M. & Bassett, D. S. The energy landscape underpinning module dynamics in the human brain connectome. NeuroImage vol. 157 364–380 (2017).

6. Yamashita A, Rothlein D, Kucyi A, Valera EM, Esterman M. Brain state-based detection of attentional fluctuations and their modulation. Neuroimage. 2021 Aug 1;236:118072.

7. Regonia PR, Takamura M, Nakano T, Ichikawa N, Fermin A, Okada G, Okamoto Y, Yamawaki S, Ikeda K, Yoshimoto J. Modeling heterogeneous brain dynamics of depression and melancholia using energy landscape analysis. Frontiers in Psychiatry. 2021 Nov 25;12:780997.

8. Xing L, Guo Z, Long Z. Energy landscape analysis of brain network dynamics in Alzheimer’s disease. Frontiers in Aging Neuroscience. 2024 May 15;16:1375091.

9. Ishida T, Yamada S, Yasuda K, Uenishi S, Tamaki A, Tabata M, Ikeda N, Takahashi S, Kimoto S. Aberrant brain dynamics of large-scale functional networks across schizophrenia and mood disorder. NeuroImage: Clinical. 2024 Jan 1;41:103574.

10. Saberi, M., Khosrowabadi, R., Khatibi, A., Misic, B. & Jafari, G. Topological impact of negative links on the stability of resting-state brain network. Scientific Reports vol. 11 (2021).

11. Heider F. Attitudes and cognitive organization. Journal of psychology. 21(1):107–12 (1946).

12. Heider F. The psychology of interpersonal relations. Psychology Press; 2013 May 13.

13. Marvel SA, Strogatz SH, Kleinberg JM. Energy landscape of social balance. Physical review letters. 2009 Nov 6;103(19):198701.

14. Kirkley A, Cantwell GT, Newman ME. Balance in signed networks. Physical Review E. 2019 Jan;99(1):012320.

15. Antal T, Krapivsky PL, Redner S. Dynamics of social balance on networks. Physical Review E—Statistical, Nonlinear, and Soft Matter Physics. 2005 Sep;72(3):036121.

16. Marvel SA, Kleinberg J, Kleinberg RD, Strogatz SH. Continuous-time model of structural balance. Proceedings of the National Academy of Sciences. 2011 Feb 1;108(5):1771–6.

17. Shojaei R, Manshour P, Montakhab A. Phase transition in a network model of social balance with Glauber dynamics. Physical Review E. 2019 Aug;100(2):022303.

18. Noudehi MG, Kargaran A, Azimi-Tafreshi N, Jafari GR. Second-to first-order phase transition: Coevolutionary versus structural balance. Physical Review E. 2022 Oct;106(4):044303.

19. Facchetti G, Iacono G, Altafini C. Computing global structural balance in large-scale signed social networks. Proceedings of the National Academy of Sciences. 2011 Dec 27;108(52):20953–8.

20. Saberi, M., Khosrowabadi, R., Khatibi, A., Misic, B. & Jafari, G. Pattern of frustration formation in the functional brain network. Network Neuroscience vol. 6 1334–1356 (2022).

21. Saberi, M., Khosrowabadi, R., Khatibi, A., Misic, B. & Jafari, G. Requirement to change of functional brain network across the lifespan. PLOS ONE vol. 16 e0260091 (2021).

22. Talesh, A. et al. Balance-energy of resting state network in obsessive-compulsive disorder. Scientific Reports vol. 13 (2023).

23. Moradimanesh, Z., Khosrowabadi, R., Eshaghi Gordji, M. & Jafari, G. R. Altered structural balance of resting-state networks in autism. Scientific Reports vol. 11 (2021).

24. Fakhari, R., Moradi, A., Ebrahimpour, R., & Khosrowabadi, R. Structural balance of resting-state brain network in Attention Deficit Hyperactivity Disorder. Basic and Clinical Neuroscience Journal (2024).

25. Soleymani, F., Khosrowabadi, R., Pedram, M. M. & Hatami, J. Impact of negative links on the structural balance of brain functional network during emotion processing. Scientific Reports vol. 13 (2023).

26. Kashyap, R. et al. The perturbational map of low frequency repetitive transcranial magnetic stimulation of primary motor cortex in movement disorders. Brain Disorders vol. 9 100071 (2023).

27. Power, J. D. et al. Functional Network Organization of the Human Brain. Neuron vol. 72 665–678 (2011).

28. Schaefer A, Kong R, Gordon EM, Laumann TO, Zuo XN, Holmes AJ, Eickhoff SB, Yeo BT. Local-global parcellation of the human cerebral cortex from intrinsic functional connectivity MRI. Cerebral cortex. 2018 Sep 1;28(9):3095–114.

29. Ezaki, T., Watanabe, T., Ohzeki, M. & Masuda, N. Energy landscape analysis of neuroimaging data. Philosophical Transactions of the Royal Society A: Mathematical, Physical and Engineering Sciences vol. 375 20160287 (2017).

30. Preti MG, Bolton TA, Van De Ville D. The dynamic functional connectome: State-of-the-art and perspectives. Neuroimage. 2017 Oct 15;160:41–54.

31. Quinn AJ, Vidaurre D, Abeysuriya R, Becker R, Nobre AC, Woolrich MW. Task-evoked dynamic network analysis through hidden Markov modeling. Frontiers in neuroscience. 2018 Aug 28;12:603.

32. Sporns O, Kötter R. Motifs in brain networks. PLoS biology. 2004 Nov;2(11):e369.

33. Wei Y, Liao X, Yan C, He Y, Xia M. Identifying topological motif patterns of human brain functional networks. Human brain mapping. 2017 May;38(5):2734–50.

34. Duclos C, Nadin D, Mahdid Y, Tarnal V, Picton P, Vanini G, Golmirzaie G, Janke E, Avidan MS, Kelz MB, Mashour GA. Brain network motifs are markers of loss and recovery of consciousness. Scientific reports. 2021 Feb 16;11(1):3892.

35. Barbey, A. K., Koenigs, M. & Grafman, J. Dorsolateral prefrontal contributions to human working memory. Cortex vol. 49 1195–1205 (2013).

36. Smith, E. E. & Jonides, J. Storage and Executive Processes in the Frontal Lobes. Science vol. 283 1657–1661 (1999).

37. Curtis, C. E. & D’Esposito, M. Persistent activity in the prefrontal cortex during working memory. Trends in Cognitive Sciences vol. 7 415–423 (2003).

38. Collette, F., Hogge, M., Salmon, E. & Van der Linden, M. Exploration of the neural substrates of executive functioning by functional neuroimaging. Neuroscience vol. 139 209–221 (2006).

39. Jung, W. H., Lee, T. Y., Yoon, Y. B., Choi, C.-H. & Kwon, J. S. Beyond Domain-Specific Expertise: Neural Signatures of Face and Spatial Working Memory in Baduk (Go Game) Experts. Frontiers in Human Neuroscience vol. 12 (2018).

40. Rosenberg, M. D. et al. Behavioral and Neural Signatures of Working Memory in Childhood. The Journal of Neuroscience vol. 40 5090–5104 (2020).

41. Chamberlain, S. R. & Sahakian, B. J. The neuropsychiatry of impulsivity. Current Opinion in Psychiatry vol. 20 255–261 (2007).

42. Penadés, R. et al. Impaired response inhibition in obsessive compulsive disorder. European Psychiatry vol. 22 404–410 (2006).

43. Robbins, T. W. Shifting and stopping: fronto-striatal substrates, neurochemical modulation and clinical implications. Philosophical Transactions of the Royal Society B: Biological Sciences vol. 362 917–932 (2007).

44. Rubia, K. et al. Progressive increase of frontostriatal brain activation from childhood to adulthood during event-related tasks of cognitive control. Human Brain Mapping vol. 27 973–993 (2006).

45. Eagle, D. M. et al. Stop-Signal Reaction-Time Task Performance: Role of Prefrontal Cortex and Subthalamic Nucleus. Cerebral Cortex vol. 18 178–188 (2007).

46. Rubia, K. et al. Mapping Motor Inhibition: Conjunctive Brain Activations across Different Versions of Go/No-Go and Stop Tasks. NeuroImage vol. 13 250–261 (2001).

47. Aron, A. R., Robbins, T. W. & Poldrack, R. A. Inhibition and the right inferior frontal cortex. Trends in Cognitive Sciences vol. 8 170–177 (2004).

48. Rieger, M., Gauggel, S. & Burmeister, K. Inhibition of ongoing responses following frontal, nonfrontal, and basal ganglia lesions. Neuropsychology vol. 17 272–282 (2003).

49. Eagle, D. M. & Robbins, T. W. Inhibitory Control in Rats Performing a Stop-Signal Reaction-Time Task: Effects of Lesions of the Medial Striatum and d-Amphetamine. Behavioral Neuroscience vol. 117 1302–1317 (2003).

50. Van Den Wildenberg, W. P. M., et al. Stimulation of the Subthalamic Region Facilitates the Selection and Inhibition of Motor Responses in Parkinson’s Disease. Journal of Cognitive Neuroscience vol. 18 626–636 (2006).

51. Bonnelle, V. et al. Salience network integrity predicts default mode network function after traumatic brain injury. Proceedings of the National Academy of Sciences vol. 109 4690–4695 (2012).

52. Leber, A. B., Turk-Browne, N. B. & Chun, M. M. Neural predictors of moment-to-moment fluctuations in cognitive flexibility. Proceedings of the National Academy of Sciences vol. 105 13592–13597 (2008).

53. Kim, C., Johnson, N. F., Cilles, S. E. & Gold, B. T. Common and Distinct Mechanisms of Cognitive Flexibility in Prefrontal Cortex. The Journal of Neuroscience vol. 31 4771–4779 (2011).

54. Rikhye, R. V., Gilra, A. & Halassa, M. M. Thalamic regulation of switching between cortical representations enables cognitive flexibility. Nature Neuroscience vol. 21 1753–1763 (2018).

55. Miyake, A. et al. The Unity and Diversity of Executive Functions and Their Contributions to Complex “Frontal Lobe” Tasks: A Latent Variable Analysis. Cognitive Psychology vol. 41 49–100 (2000).

56. Yehene, E., Meiran, N. & Soroker, N. Basal Ganglia Play a Unique Role in Task Switching within the Frontal-Subcortical Circuits: Evidence from Patients with Focal Lesions. Journal of Cognitive Neuroscience vol. 20 1079–1093 (2008).

57. Kincade, J. M., Abrams, R. A., Astafiev, S. V., Shulman, G. L. & Corbetta, M. An Event-Related Functional Magnetic Resonance Imaging Study of Voluntary and Stimulus-Driven Orienting of Attention. The Journal of Neuroscience vol. 25 4593– 4604 (2005).

58. Corbetta, M. & Shulman, G. L. Control of goal-directed and stimulus-driven attention in the brain. Nature Reviews Neuroscience vol. 3 201–215 (2002).

59. Vossel, S., Geng, J. J. & Fink, G. R. Dorsal and Ventral Attention Systems. The Neuroscientist vol. 20 150–159 (2013).

60. Tamber-Rosenau, B. J., Asplund, C. L. & Marois, R. Functional dissociation of the inferior frontal junction from the dorsal attention network in top-down attentional control. Journal of Neurophysiology vol. 120 2498–2512 (2018).

61. Wallis, G., Stokes, M., Cousijn, H., Woolrich, M. & Nobre, A. C. Frontoparietal and Cingulo-opercular Networks Play Dissociable Roles in Control of Working Memory. Journal of Cognitive Neuroscience vol. 27 2019–2034 (2015).

62. Cao, H. & Cannon, T. D. Distinct and temporally associated neural mechanisms underlying concurrent, postsuccess, and posterror cognitive controls: Evidence from a stop-signal task. Human Brain Mapping vol. 42 2677–2690 (2021).

63. Yin, S., Deák, G. & Chen, A. Coactivation of cognitive control networks during task switching. Neuropsychology vol. 32 31–39 (2018).

64. He, X. et al. Abnormal salience network in normal aging and in amnestic mild cognitive impairment and Alzheimer’s disease. Human Brain Mapping vol. 35 3446–3464 (2013).

65. Onoda, K., Ishihara, M. & Yamaguchi, S. Decreased Functional Connectivity by Aging Is Associated with Cognitive Decline. Journal of Cognitive Neuroscience vol. 24 2186–2198 (2012).

66. Ruiz-Rizzo, A. L. et al. Decreased cingulo-opercular network functional connectivity mediates the impact of aging on visual processing speed. Neurobiology of Aging vol. 73 50–60 (2019).

67. Hausman, H. K. et al. The Role of Resting-State Network Functional Connectivity in Cognitive Aging. Frontiers in Aging Neuroscience vol. 12 (2020).

68. Geerligs, L., Renken, R. J., Saliasi, E., Maurits, N. M. & Lorist, M. M. A Brain-Wide Study of Age-Related Changes in Functional Connectivity. Cerebral Cortex vol. 25 1987–1999 (2014).

69. Richiardi, J., Achard, S., Bunke, H. & Van De Ville, D. Machine Learning with Brain Graphs: Predictive Modeling Approaches for Functional Imaging in Systems Neuroscience. IEEE Signal Processing Magazine vol. 30 58–70 (2013).

70. Kazeminejad, A., Golbabaei, S. & Soltanian-Zadeh, H. Graph theoretical metrics and machine learning for diagnosis of Parkinson’s disease using rs-fMRI. 2017 Artificial Intelligence and Signal Processing Conference (AISP) (2017) doi:10.1109/aisp.2017.8324124.

71. Tolan, E. & Isik, Z. Graph Theory Based Classification of Brain Connectivity Network for Autism Spectrum Disorder. Bioinformatics and Biomedical Engineering 520–530 (2018) doi:10.1007/978-3-319-78723-7_45.

72. Khazaee, A., Ebrahimzadeh, A. & Babajani-Feremi, A. Application of advanced machine learning methods on resting-state fMRI network for identification of mild cognitive impairment and Alzheimer’s disease. Brain Imaging and Behavior vol. 10 799–817 (2015).

73. Kılıç, B. & Aydın, S. Classification of Contrasting Discrete Emotional States Indicated by EEG Based Graph Theoretical Network Measures. Neuroinformatics vol. 20 863–877 (2022).

74. Garcia-Ramos, C. et al. Network phenotypes and their clinical significance in temporal lobe epilepsy using machine learning applications to morphological and functional graph theory metrics. Scientific Reports vol. 12 (2022).

75. Liu, W. et al. Graph-Theory-Based Degree Centrality Combined with Machine Learning Algorithms Can Predict Response to Treatment with Antipsychotic Medications in Patients with First-Episode Schizophrenia. Disease Markers vol. 2022 1–7 (2022).

76. Rieck, J. R., Baracchini, G., Nichol, D., Abdi, H. & Grady, C. L. Dataset of functional connectivity during cognitive control for an adult lifespan sample. Data in Brief vol. 39 107573 (2021).

77. R Core Team. R: A language and environment for statistical computing. R Foundation for Statistical Computing, Vienna, Austria (2023). URL https://www.R-project.org/

78. Wickham, H. ggplot2: Elegant Graphics for Data Analysis (Springer-Verlag New York, 2016). URL https://ggplot2.tidyverse.org

79. Csardi, G. & Nepusz, T. The igraph software package for complex network research. InterJournal, Complex Systems 1695 (2006). URL http://igraph.org

80. Watson, C. G. brainGraph: Graph Theory Analysis of Brain MRI Data. R package version 3.0.0 (2020). URL https://CRAN.R-project.org/package=brainGraph

81. Renedo Mirambell, M. clustAnalytics: Cluster Evaluation on Graphs (R package version 0.5.4) [Computer software] (2023). URL https://CRAN.R-project.org/package=clustAnalytics

82. Kuhn, M. Caret: Classification and Regression Training. R package version (2008). URL https://CRAN.R-project.org/package=caret

83. Meyer, D., et al. e1071: Misc Functions of the Department of Statistics, Probability Theory Group (formerly: E1071), TU Wien. R package version 1.2, (2019). Available at https://CRAN.R-project.org/package=e1071 (Accessed: date of access).

