## Supplementary Material for "The brain selectively allocates energy to functional brain networks under cognitive control"

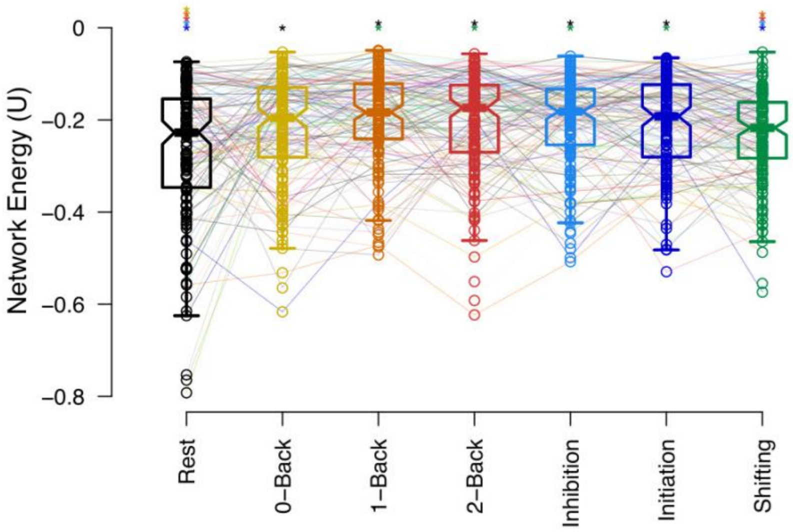


**Figure S1. Energy of whole-brain network during cognitive control tasks and resting-state extracted based on Schaefer's atlas.** Horizontal lines represent group-level medians, while notches on the boxes signify the 95% confidence interval for the median. Individual subjects are linked by lines, with task conditions differentiated by color. Colored asterisks above their respective conditions mark significant corrected p-values for non-parametric pair-wise comparisons between cognitive task conditions.

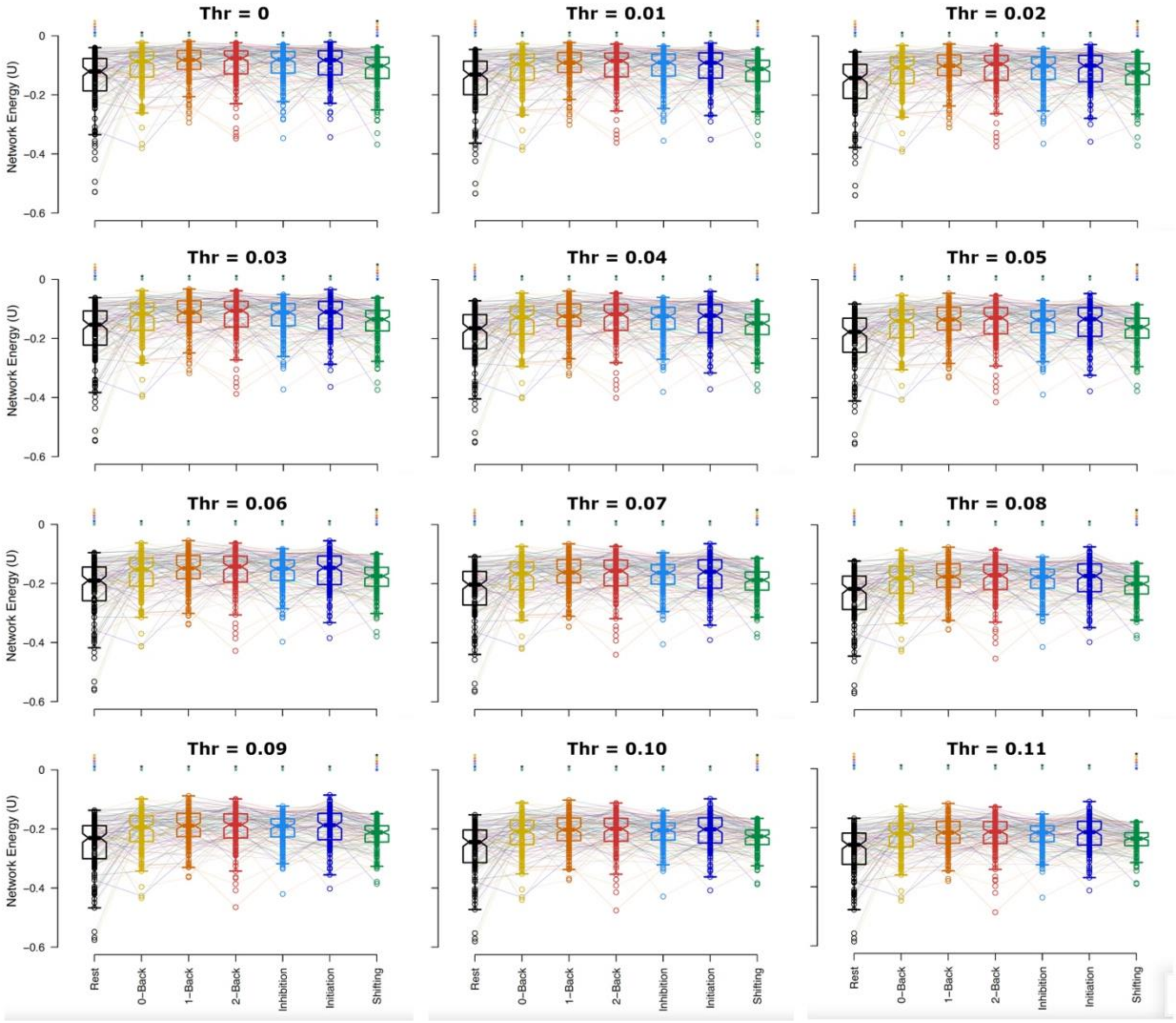


**Figure S2.** Effect of thresholding on the energy level of the whole-brain network during cognitive control tasks and resting-state, extracted using Power's atlas. Thresholds were applied to positive and negative connections with absolute correlation values less than the indicated thresholds before computing network energy. Horizontal lines represent group-level medians, and notches on the boxes indicate the 95% confidence intervals for the medians. Individual subjects are connected by lines, with task conditions distinguished by color. Colored asterisks above the respective conditions denote significant corrected p-values for non-parametric pairwise comparisons between cognitive task conditions.

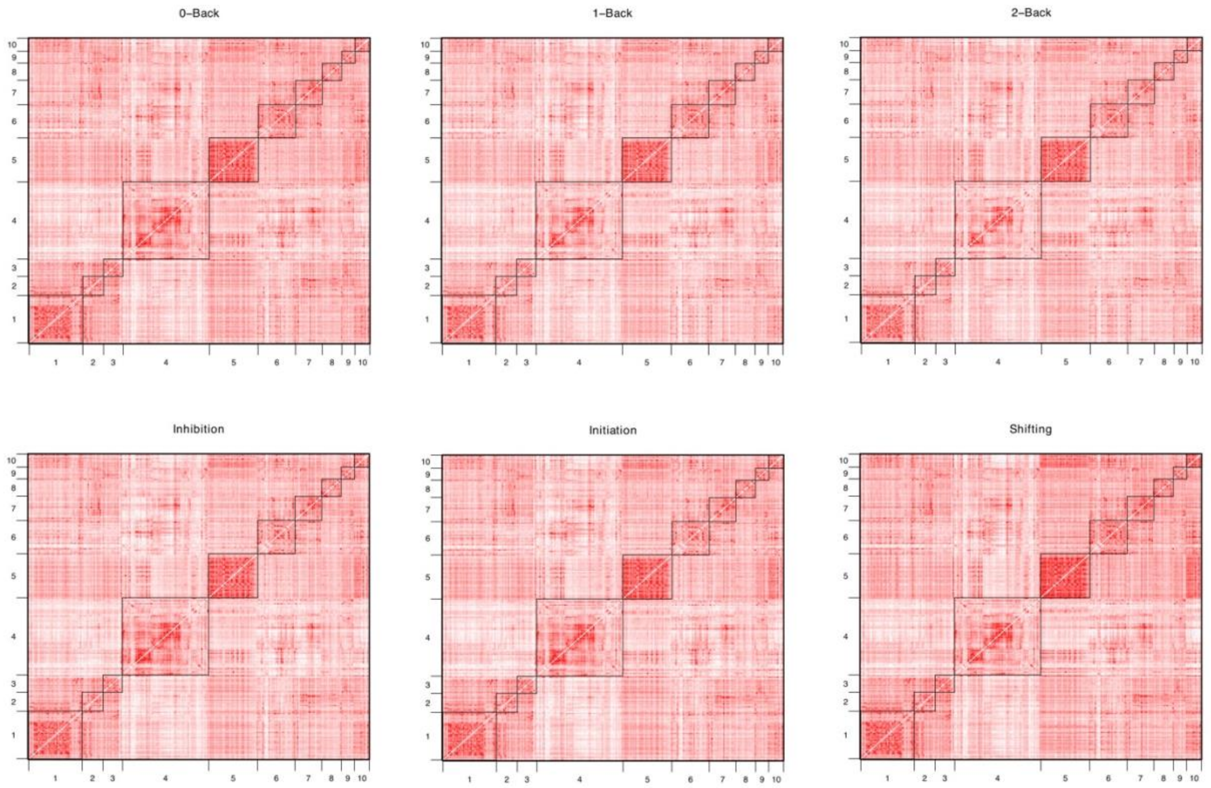


**Figure S3.** Group-average functional connectivity across all subjects for each task condition. The 229 Power's regions of interest were categorized into 10 canonical functional networks. Internetwork connections are represented by squares, with the intensity of colors indicating the strength of the group-average connections. Network labels are as follows: 1: Somatosensory, 2: Cingulate-Opercular, 3: Auditory, 4: Default Mode, 5: Visual, 6: Prefrontal, 7: Salience, 8: Subcortical, 9: Ventral Attention, 10: Dorsal Attention.

| | Rest | 0-Back | 1-Back | 2-Back | Inhibition | Initiation | Shifting |
|---|---|---|---|---|---|---|---|
| **Rest** | - | - | - | - | - | - | - |
| **0-Back** | 0.018 (5.7E-5) | - | - | - | - | - | - |
| **1-Back** | 0.033 (2.93E-7) | 0.011 (0.370) | - | - | - | - | |
| **2-Back** | 0.348 (3.48E-7) | 0.004 (0.370) | 0.001 (0.987) | - | - | - | - |
| **Inhibition** | 0.031 (4.54E-7) | 0.005 (0.525) | 0.001 (0.820) | -0.001 (0.820) | - | - | - |
| **Initiation** | 0.035 (6.22E-7) | 0.001 (0.525) | -0.001 (0.808) | -0.005 (0.808) | -0.001 (0.987) | - | - |
| **Shifting** | 0.013 (0.040) | -0.018 (0.031) | -0.020 (5.51E-4) | -0.015 (8.82 E-4) | -0.013 (0.001) | -0.025 (0.001) | - |

**Table S1. Pairwise Comparisons of Whole-Brain Network Energy Levels Across Various Cognitive Task Conditions.** Cells indicate the median of paired differences among subjects, with p-values (in parentheses) from the Wilcoxon signed-rank test after applying multiple comparison correction. Highlighted cells represent comparisons with a corrected p-value less than 0.05.

| | AUD | COP | DAN | DMN | FPC | SAL | SM | SC | VAN | VIS |
|---|---|---|---|---|---|---|---|---|---|---|
| **AUD** | - | - | - | - | - | - | - | - | - | - |
| **COP** | -0.046 (0.002) | - | - | - | - | - | - | - | - | - |
| **DAN** | -0.037 (0.326) | 0.027 (0.067) | - | - | - | - | - | - | - | - |
| **DMN** | 0.079 (4.19E-8) | 0.129 (8.81E-18) | 0.119 (1.45E-10) | - | - | - | - | - | - | - |
| **FPC** | 0.072 (3.95E-6) | 0.129 (2.27E-14) | 0.097 (2.96E-8) | -0.010 (0.444) | - | - | - | - | - | - |
| **SAL** | 0.030 (0.047) | 0.079 (7.32E-8) | 0.051 (0.002) | -0.051 (6.38E-5) | -0.040 (0.002) | - | - | - | - | - |
| **SM** | -0.052 (0.583) | -0.015 (0.001) | -0.029 (0.461) | -0.144 (1.38E-10) | -0.128 (2.96E-8) | -0.085 (0.008) | - | - | - | - |
| **SC** | -0.002 (0.014) | 0.042 (0.958) | 0.015 (0.141) | -0.084 (4.05E-12) | -0.081 (4.73E-10) | -0.040 (3.82E-5) | 0.037 (0.014) | - | - | - |
| **VAN** | 0.006 (0.434) | 0.067 (2.28E-4) | 0.054 (0.057) | -0.083 (3.82E-5) | -0.065 (9.39E-4) | -0.014 (0.423) | 0.073 (0.233) | 0.014 (0.002) | - | - |
| **VIS** | -0.108 (6.65E-10) | -0.057 (8.18E-4) | -0.080 (1.12E-6) | -0.197 (1.46E-28) | -0.183 (2.81E-24) | -0.148 (4.21E-17) | -0.047 (3.79E-11) | -0.108 (0.009) | -0.136 (1.19E-10) | - |

**Table S2. Pairwise Comparisons of Canonical Networks' Energies During Resting State.** Cells indicate the median of paired differences among subjects, with p-values (in parentheses) from the Wilcoxon signed-rank test after applying multiple comparison correction. Highlighted cells represent comparisons with a corrected p-value less than 0.05.

| | AUD | COP | DAN | DMN | FPC | SAL | SM | SC | VAN | VIS |
|---|---|---|---|---|---|---|---|---|---|---|
| **AUD** | - | - | - | - | - | - | - | - | - | - |
| **COP** | -0.015 (0.679) | - | - | - | - | - | - | - | - | - |
| **DAN** | -0.018 (0.252) | 0.002 (0.679) | - | - | - | - | - | - | - | - |
| **DMN** | 0.031 (8.68E-4) | 0.05 (3.96E-5) | 0.057 (2.41E-7) | - | - | - | - | - | - | - |
| **FPC** | -0.014 (0.776) | 0.01 (0.846) | 0.013 (0.465) | -0.045 (3.96E-5) | - | - | | - | - | - |
| **SAL** | -0.003 (0.776) | 0.004 (0.846) | 0.012 (0.514) | -0.035 (8.71E-5) | -0.012 (0.944) | - | - | - | - | - |
| **SM** | -0.015 (0.498) | -0.005 (0.846) | -0.01 (0.846) | -0.059 (1.88E-5) | -0.012 (0.679) | -0.016 (0.679) | - | - | - | - |
| **SC** | -0.013 (0.114) | 0.005 (0.558) | -0.003 (0.944) | -0.049 (2.12E-8) | -0.028 (0.373) | -0.02 (0.418) | 0.000615 (0.775) | - | - | - |
| **VAN** | -0.008 (0.776) | 0.01 (0.846) | 0.016 (0.511) | -0.047 (2.34E-4) | -0.005 (0.944) | 0.004 (0.944) | 0.010308 (0.688) | 0.008 (0.498) | - | - |
| **VIS** | -0.09 (3.44E-10) | -0.079 (9.13E-8) | -0.078 (2.31E-7) | -0.132 (1.31E-22) | -0.08 (9.33E-10) | -0.07585 (3.69E-9) | -0.067 (4.77E-7) | -0.074 (2.31E-7) | -0.076 (4.50E-8) | - |

**Table S3. Pairwise Comparisons of Canonical Networks' Energies During 0-Back Task.** Cells indicate the median of paired differences among subjects, with p-values (in parentheses) from the Wilcoxon signed-rank test after applying multiple comparison correction. Highlighted cells represent comparisons with a corrected p-value less than 0.05.

| | AUD | COP | DAN | DMN | FPC | SAL | SM | SC | VAN | VIS |
|---|---|---|---|---|---|---|---|---|---|---|
| **AUD** | - | - | - | - | - | - | - | - | - | - |
| **COP** | 0.001 (0.858) | - | - | - | - | - | - | - | - | - |
| **DAN** | -0.019 (0.105) | -0.016 (0.154) | - | - | - | - | - | - | - | - |
| **DMN** | 0.029 (0.001) | 0.034 (0.001) | 0.057 (1.17E-07) | - | - | - | - | - | - | - |
| **FPC** | -0.01 (0.150) | -0.012 (0.217) | 0.004 (0.827) | -0.043 (4.73E-8) | - | - | - | - | - | - |
| **SAL** | -0.019 (0.150) | -0.013 (0.219) | -0.003 (0.827) | -0.052 (1.34E-7) | -0.006 (0.998) | - | - | - | - | - |
| **SM** | -0.028 (0.290) | -0.023 (0.319) | -0.002 (0.646) | -0.059 (1.93E-06) | -0.024 (0.858) | -0.007 (0.858) | - | - | - | - |
| **SC** | -0.013 (0.009) | -0.015 (0.017) | 0.003 (0.563) | -0.041 (1.14E-10) | -0.005 (0.361) | 0.001 (0.395) | 0.015 (0.315) | - | - | - |
| **VAN** | -0.006 (0.467) | -0.011 (0.563) | 0.008 (0.467) | -0.051 (3.89E-05) | 0.001 (0.563) | 0.002 (0.570) | 0.016 (0.784) | -0.002 (0.150) | - | - |
| **VIS** | -0.086 (2.69E-11) | -0.077 (1.09E-10) | -0.06 (4.11E-07) | -0.131 (2.69E-24) | -0.08 (5.97E-09) | -0.07 (2.64E-8) | -0.059 (2.64E-8) | -0.08 (3.27E-6) | -0.074 (2.64E-8) | - |

**Table S4. Pairwise Comparisons of Canonical Networks' Energies During 1-Back Task.** Cells indicate the median of paired differences among subjects, with p-values (in parentheses) from the Wilcoxon signed-rank test after applying multiple comparison correction. Highlighted cells represent comparisons with a corrected p-value less than 0.05.

| | AUD | COP | DAN | DMN | FPC | SAL | SM | SC | VAN | VIS |
|---|---|---|---|---|---|---|---|---|---|---|
| **AUD** | - | - | - | - | - | - | - | - | - | - |
| **COP** | 0.005 (0.405) | - | - | - | - | - | - | - | - | - |
| **DAN** | -0.013 (0.084) | -0.029 (0.011) | - | - | - | - | - | - | - | - |
| **DMN** | 0.044 (1.22E-07) | 0.038 (5.21E-05) | 0.068 (1.05E-12) | - | - | - | - | - | - | - |
| **FPC** | -0.014 (0.340) | -0.019 (0.084) | 0.006 (0.405) | -0.055 (4.69E-11) | - | - | - | - | - | - |
| **SAL** | -0.004 (0.938) | -0.003 (0.391) | 0.012 (0.087) | -0.048 (2.51E-08) | 0.006 (0.391) | - | - | - | - | - |
| **SM** | -0.010 (0.279) | -0.018 (0.057) | -0.001 (0.710) | -0.064 (2.01E-10) | -0.007 (0.785) | -0.017 (0.292) | - | - | - | - |
| **SC** | -0.017 (0.094) | -0.023 (0.013) | 0.001 (0.949) | -0.073 (1.93E-12) | -0.018 (0.441) | -0.023 (0.104) | 0.004 (0.726) | - | - | - |
| **VAN** | -0.002 (0.938) | -0.022 (0.405) | 0.013 (0.173) | -0.053 (8.20E-07) | 0.007 (0.456) | -0.008 (0.984) | 0.015 (0.391) | 0.015 (0.217) | - | - |
| **VIS** | -0.088 (8.77E-13) | -0.095 (7.13E-14) | -0.071 (4.71E-09) | -0.140 (1.24E-28) | -0.075 (1.31E-11) | -0.089 (3.50E-13) | -0.064 (5.92E-09) | -0.062 (4.5E-09) | -0.087 (4.00E-10) | - |

**Table S5. Pairwise Comparisons of Canonical Networks' Energies During 2-Back Task.** Cells indicate the median of paired differences among subjects, with p-values (in parentheses) from the Wilcoxon signed-rank test after applying multiple comparison correction. Highlighted cells represent comparisons with a corrected p-value less than 0.05.

| | AUD | COP | DAN | DMN | FPC | SAL | SM | SC | VAN | VIS |
|---|---|---|---|---|---|---|---|---|---|---|
| **AUD** | - | - | - | - | - | - | - | - | - | - |
| **COP** | -0.012 (0.533) | - | - | - | - | - | - | - | - | - |
| **DAN** | -0.021 (0.111) | -0.006 (0.461) | - | - | - | - | - | - | - | - |
| **DMN** | 0.031 (8.84E-4) | 0.047 (1.36E-05) | 0.057 (4.97E-08) | - | - | - | - | - | - | - |
| **FPC** | 0.029 (0.046) | 0.037 (0.002) | 0.044 (2.93E-05) | -0.013 (0.229) | - | - | - | - | - | - |
| **SAL** | -0.004 (0.515) | 0.002 (0.992) | 0.010 (0.366) | -0.056 (5.08E-06) | -0.029 (0.001) | - | - | - | - | - |
| **SM** | -0.031 (0.218) | -0.015 (0.583) | -0.001 (0.749) | -0.062 (3.29E-07) | -0.054 (1.18E-4) | -0.015 (0.533) | - | - | - | - |
| **SC** | -0.025 (0.020) | -0.006 (0.156) | -0.011 (0.704) | -0.064 (3.39E-10) | -0.048 (8.61E-07) | -0.018 (0.156) | 0.001 (0.488) | - | - | - |
| **VAN** | -0.009 (0.747) | 0.007 (0.813) | 0.002 (0.322) | -0.046 (9.32E-05) | -0.029 (0.008) | 0.0003 (0.749) | 0.024 (0.420) | 0.012 (0.097) | - | - |
| **VIS** | -0.088 (2.60E-09) | -0.070 (6.83E-08) | -0.059 (1.16E-06) | -0.125 (4.87E-23) | -0.114 (1.96E-18) | -0.068 (1.03E-08) | -0.053 (4.09E-07) | -0.057 (4.96E-06) | -0.073 (4.32E-08) | - |

**Table S6. Pairwise Comparisons of Canonical Networks' Energies During Inhibition Condition of Go/No-go Task.** Cells indicate the median of paired differences among subjects, with p-values (in parentheses) from the Wilcoxon signed-rank test after applying multiple comparison correction. Highlighted cells represent comparisons with a corrected p-value less than 0.05.

| | AUD | COP | DAN | DMN | FPC | SAL | SM | SC | VAN | VIS |
|---|---|---|---|---|---|---|---|---|---|---|
| **AUD** | - | - | - | - | - | - | - | - | - | - |
| **COP** | -0.007<br>(0.384) | - | - | - | - | - | - | - | - | - |
| **DAN** | -0.012<br>(0.367) | -0.003<br>0.929 | - | - | - | - | - | - | - | - |
| **DMN** | 0.033<br>(5.51E-05) | 0.044<br>(8.02E-07) | 0.058<br>(1.74E-07) | - | - | - | - | - | - | - |
| **FPC** | 0.025<br>(0.024) | 0.036<br>(8.29E-04) | 0.037<br>(5.58E-04) | -0.014<br>(0.155) | - | - | - | - | - | - |
| **SAL** | -0.004<br>(0.926) | 0.012<br>(0.517) | 0.005<br>(0.537) | -0.040<br>(4.43E-05) | -0.020<br>(0.016) | - | - | - | - | - |
| **SM** | -0.0366<br>(0.537) | -0.011<br>(0.826) | -0.013<br>(0.778) | -0.06896<br>(6.53E-06) | -0.06135<br>(0.003) | -0.036<br>(0.638) | - | - | - | - |
| **SC** | -0.025<br>(0.021) | 0.009<br>(0.289) | 0.007<br>(0.232) | -0.055<br>(2.90E-11) | -0.032<br>(8.02E-07) | -0.018<br>(0.044) | 0.012<br>(0.122) | - | - | - |
| **VAN** | 0.002<br>(0.617) | 0.007<br>(0.848) | 0.014<br>(0.860) | -0.047<br>(8.77E-06) | -0.026<br>(0.004) | -0.013<br>(0.623) | 0.028<br>(0.929) | 0.008<br>(0.204) | - | - |
| **VIS** | -0.085<br>(6.13E-10) | -0.077<br>(4.16E-07) | -0.076<br>(1.34E-07) | -0.129<br>(2.96E-23) | -0.121<br>(5.57E-17) | -0.081<br>(1.87E-09) | -0.042<br>(1.63E-07) | -0.073<br>(2.41E-05) | -0.091<br>(1.63E-07) | - |

**Table S7. Pairwise Comparisons of Canonical Networks' Energies During Initiation Condition of Go/No-go Task.** Cells indicate the median of paired differences among subjects, with p-values (in parentheses) from the Wilcoxon signed-rank test after applying multiple comparison correction. Highlighted cells represent comparisons with a corrected p-value less than 0.05.

| | AUD | COP | DAN | DMN | FPC | SAL | SM | SC | VAN | VIS |
|---|---|---|---|---|---|---|---|---|---|---|
| **AUD** | - | - | - | - | - | - | - | - | - | - |
| **COP** | -0.006<br>(0.261) | - | - | - | - | - | - | - | - | - |
| **DAN** | -0.079<br>(7.47E-08) | -0.058<br>(1.74E-05) | - | - | - | - | - | - | - | - |
| **DMN** | 0.067<br>(1.82E-11) | 0.094<br>(2.29E-15) | 0.154<br>(1.76E-30) | - | - | - | - | - | - | - |
| **FPC** | 0.005<br>(0.736) | 0.015<br>(0.145) | 0.077<br>(1.76E-10) | -0.071<br>(1.77E-12) | - | - | - | - | - | - |
| **SAL** | -0.005<br>(0.679) | 0.018<br>(0.408) | 0.059<br>(3.48E-08) | -0.085<br>(2.21E-14) | -0.013<br>(0.489) | - | - | - | - | - |
| **SM** | -0.027<br>(0.009) | 0.001<br>(0.153) | 0.056<br>(0.002) | -0.092<br>(3.13E-21) | -0.022<br>(0.0008) | -0.004<br>(0.010) | - | - | - | - |
| **SC** | -0.042<br>(0.170) | -0.009<br>(0.894) | 0.031<br>(1.24E-06) | -0.105<br>(5.37E-19) | -0.033<br>(0.099) | -0.031<br>(0.325) | -0.021<br>(0.100) | - | - | - |
| **VAN** | 0.005<br>(0.357) | 0.034<br>(0.035) | 0.075<br>(1.41E-10) | -0.066<br>(5.50E-09) | 0.005<br>(0.446) | 0.006<br>(0.157) | 0.031<br>(1.87E-04) | 0.047<br>(0.014) | - | - |
| **VIS** | -0.122<br>(5.37E-19) | -0.111<br>(1.34E-15) | -0.059<br>(2.77E-06) | -0.211<br>(4.91E-39) | -0.121<br>(8.14E-24) | -0.132<br>(6.14E-20) | -0.115<br>(1.87E-13) | -0.099<br>(8.23E-19) | -0.140<br>(2.24E-21) | - |

**Table S8. Pairwise Comparisons of Canonical Networks' Energies During Shifting Task.** Cells indicate the median of paired differences among subjects, with p-values (in parentheses) from the Wilcoxon signed-rank test after applying multiple comparison correction. Highlighted cells represent comparisons with a corrected p-value less than 0.05.

| | Rest | 0-Back | 1-Back | 2-Back | Inhibition | Initiation | Shifting |
|---|---|---|---|---|---|---|---|
| **Rest** | - | - | - | - | - | - | - |
| **0-Back** | 0.018<br>(5.7E-5) | - | - | - | - | - | - |
| **1-Back** | 0.033<br>(2.93E-7) | 0.011<br>(0.370) | - | - | - | - | |
| **2-Back** | 0.348<br>(3.48E-7) | 0.004<br>(0.370) | 0.001<br>(0.987) | - | - | - | - |
| **Inhibition** | 0.031<br>(4.54E-7) | 0.005<br>(0.525) | 0.001<br>(0.820) | -0.001<br>(0.820) | - | - | - |
| **Initiation** | 0.035<br>(6.22E-7) | 0.001<br>(0.525) | -0.001<br>(0.808) | -0.005<br>(0.808) | -0.001<br>(0.987) | - | - |
| **Shifting** | 0.013<br>(0.040) | -0.018<br>(0.031) | -0.020<br>(5.51E-4) | -0.015<br>(8.82 E-4) | -0.013<br>(0.001) | -0.025<br>(0.001) | - |

**Table S9. Pairwise Comparisons of Auditory Network's Energy Across Cognitive Task Conditions.** Cells indicate the median of paired differences among subjects, with p-values (in parentheses) from the Wilcoxon signed-rank test after applying multiple comparison correction. Highlighted cells represent comparisons with a corrected p-value less than 0.05.

| | Rest | 0-Back | 1-Back | 2-Back | Inhibition | Initiation | Shifting |
|---|---|---|---|---|---|---|---|
| **Rest** | - | - | - | - | - | - | - |
| **0-Back** | 0.150 (3.21E-19) | - | - | - | - | - | - |
| **1-Back** | 0.185 (4.26E-26) | 0.037 (0.006) | - | - | - | - | |
| **2-Back** | 0.190 (4.18E-28) | 0.044 (0.001) | 0.010 (0.631) | - | - | - | - |
| **Inhibition** | 0.159 (3.38E-20) | -0.001 (0.874) | -0.024 (0.002) | -0.040 (3.80E-04) | - | - | - |
| **Initiation** | 0.167 (3.38E-20) | 0.006 (0.874) | -0.018 (0.005) | -0.044 (6.48E-04) | -0.001 (0.874) | - | - |
| **Shifting** | 0.116 (4.15E-13) | -0.035 (0.002) | -0.061 (1.34E-08) | -0.080 (1.54E-10) | -0.033 (0.002) | -0.037 (0.002) | - |

**Table S10. Pairwise Comparisons of Cingulo-Opercular Network's Energy Across Cognitive Task Conditions.** Cells indicate the median of paired differences among subjects, with p-values (in parentheses) from the Wilcoxon signed-rank test after applying multiple comparison correction. Highlighted cells represent comparisons with a corrected p-value less than 0.05.

| | Rest | 0-Back | 1-Back | 2-Back | Inhibition | Initiation | Shifting |
|---|---|---|---|---|---|---|---|
| **Rest** | - | - | - | - | - | - | - |
| **0-Back** | 0.117 (2.17E-12) | - | - | - | - | - | - |
| **1-Back** | 0.128 (1.73E-15) | 0.018 (0.117) | - | - | - | - | |
| **2-Back** | 0.121 (4.85E-17) | 0.016 (0.130) | -0.001 0.803) | - | - | - | - |
| **Inhibition** | 0.114 (7.38E-12) | -0.008 (0.643) | -0.027 (0.036) | -0.029 (0.030) | - | - | - |
| **Initiation** | 0.116 (8.59E-14) | 0.008 (0.623) | -0.013 (0.287) | -0.014 (0.352) | 0.009 (0.247) | - | - |
| **Shifting** | 0.012 (0.153) | -0.088 (2.13E-11) | -0.099 (6.38E-15) | -0.112 (4.31E-17) | -0.080 (3.90E-11) | -0.107 (2.94E-13) | - |

**Table S11. Pairwise Comparisons of Dorsal Attention Network's Energy Across Cognitive Task Conditions.** Cells indicate the median of paired differences among subjects, with p-values (in parentheses) from the Wilcoxon signed-rank test after applying multiple comparison correction. Highlighted cells represent comparisons with a corrected p-value less than 0.05.

| | Rest | 0-Back | 1-Back | 2-Back | Inhibition | Initiation | Shifting |
|---|---|---|---|---|---|---|---|
| **Rest** | - | - | - | - | - | - | - |
| **0-Back** | 0.065 (8.37E-12) | - | - | - | - | - | - |
| **1-Back** | 0.086 (6.60E-19) | 0.019 (0.009) | - | - | - | - | |
| **2-Back** | 0.096 (1.06E-21) | 0.030 (1.38E-04) | 0.008 (0.183) | - | - | - | - |
| **Inhibition** | 0.065 (6.32E-11) | -0.005 (0.608) | -0.018 (0.001) | -0.029 (6.37E-06) | - | - | - |
| **Initiation** | 0.076 (1.48E-13) | 0.001 (0.587) | -0.010 (0.047) | -0.023 (8.45E-04) | 0.011 (0.286) | - | - |
| **Shifting** | 0.073 (3.15E-12) | 0.002 (0.997) | -0.026 (0.006) | -0.027 (5.13E-05) | 0.001 (0.621) | -0.007 (0.549) | - |

**Table S12. Pairwise Comparisons of Default Mode Network's Energy Across Cognitive Task Conditions.** Cells indicate the median of paired differences among subjects, with p-values (in parentheses) from the Wilcoxon signed-rank test after applying multiple comparison correction. Highlighted cells represent comparisons with a corrected p-value less than 0.05.

| | Rest | 0-Back | 1-Back | 2-Back | Inhibition | Initiation | Shifting |
|---|---|---|---|---|---|---|---|
| **Rest** | - | - | - | - | - | - | - |
| **0-Back** | 0.030 (0.004) | - | - | - | - | - | - |
| **1-Back** | 0.040 (7.09E-05) | 0.012 (0.298) | - | - | - | - | |
| **2-Back** | 0.035 (1.38E-05) | 0.012 (0.189) | 0.001 (0.752) | - | - | - | - |
| **Inhibition** | 0.056 (9.54E-08) | 0.030 (0.019) | 0.017 (0.189) | 0.012 (0.288) | - | - | - |
| **Initiation** | 0.050 (3.18E-08) | 0.027 (0.003) | 0.021 (0.059) | 0.023 (0.080) | 0.001 (0.453) | - | - |
| **Shifting** | 0.008 (0.453) | -0.013 (0.024) | -0.034 (4.55E-04) | -0.030 (6.22E-05) | -0.049 (3.02E-07) | -0.064 (9.54E-08) | - |

**Table S13. Pairwise Comparisons of Frontoparietal Network's Energy Across Cognitive Task Conditions.** Cells indicate the median of paired differences among subjects, with p-values (in parentheses) from the Wilcoxon signed-rank test after applying multiple comparison correction. Highlighted cells represent comparisons with a corrected p-value less than 0.05.

| | Rest | 0-Back | 1-Back | 2-Back | Inhibition | Initiation | Shifting |
|---|---|---|---|---|---|---|---|
| **Rest** | - | - | - | - | - | - | - |
| **0-Back** | 0.076 (2.87E-08) | - | - | - | - | - | - |
| **1-Back** | 0.087 (2.81E-11) | 0.010 (0.338) | - | - | - | - | |
| **2-Back** | 0.100 (1.74E-15) | 0.022 (0.034) | 0.007 (0.302) | - | - | - | - |
| **Inhibition** | 0.065 (4.35E-08) | -0.010 (0.585) | -0.021 (0.103) | -0.034 (0.003) | - | - | - |
| **Initiation** | 0.078 (1.11E-09) | 0.002 (0.615) | -0.010 (0.585) | -0.018 (0.076) | 0.015 (0.346) | - | - |
| **Shifting** | 0.040 (0.001) | -0.034 (0.004) | -0.048 (7.93E-05) | -0.058 (4.35E-08) | -0.030 (0.017) | -0.036 (0.002) | - |

**Table S14. Pairwise Comparisons of Salience Network's Energy Across Cognitive Task Conditions.** Cells indicate the median of paired differences among subjects, with p-values (in parentheses) from the Wilcoxon signed-rank test after applying multiple comparison correction. Highlighted cells represent comparisons with a corrected p-value less than 0.05.

| | Rest | 0-Back | 1-Back | 2-Back | Inhibition | Initiation | Shifting |
|---|---|---|---|---|---|---|---|
| **Rest** | - | - | - | - | - | - | - |
| **0-Back** | 0.114 (4.56E-14) | - | - | - | - | - | - |
| **1-Back** | 0.122 (2.07E-16) | 0.013 (0.217) | - | - | - | - | |
| **2-Back** | 0.158 (3.60E-18) | 0.025 (0.043) | 0.005 (0.573) | - | - | - | - |
| **Inhibition** | 0.119 (1.88E-12) | -0.008 (0.415) | -0.027 (0.025) | -0.025 (0.004) | - | - | - |
| **Initiation** | 0.100 (1.88E-12) | -0.009 (0.573) | -0.017 (0.053) | -0.022 (0.012) | -0.001 (0.771) | - | - |
| **Shifting** | 0.107 (2.28E-09) | -0.036 (0.004) | -0.045 (2.75E-05) | -0.056 (4.91E-07) | -0.016 (0.033) | -0.024 (0.025) | - |

**Table S15. Pairwise Comparisons of Somatomotor Network's Energy Across Cognitive Task Conditions.** Cells indicate the median of paired differences among subjects, with p-values (in parentheses) from the Wilcoxon signed-rank test after applying multiple comparison correction. Highlighted cells represent comparisons with a corrected p-value less than 0.05.

| | Rest | 0-Back | 1-Back | 2-Back | Inhibition | Initiation | Shifting |
|---|---|---|---|---|---|---|---|
| **Rest** | - | - | - | - | - | - | - |
| **0-Back** | 0.103 (2.38E-11) | - | - | - | - | - | - |
| **1-Back** | 0.121 (1.76E-17) | 0.036 (0.088) | - | - | - | - | |
| **2-Back** | 0.114 (4.50E-17) | 0.026 (0.153) | -0.010 (0.775) | - | - | - | - |
| **Inhibition** | 0.092 (1.82E-12) | -0.001 (0.666) | -0.036 (0.017) | -0.025 (0.028) | - | - | - |
| **Initiation** | 0.102 (1.99E-12) | 0.001 (0.676) | -0.024 (0.214) | -0.009 (0.333) | 0.004 (0.353) | - | - |
| **Shifting** | 0.053 (1.41E-04) | -0.059 (7.72E-05) | -0.085 (9.30E-10) | -0.075 (3.25E-09) | -0.044 (7.91E-05) | -0.057 (3.55E-06) | - |

**Table S16. Pairwise Comparisons of Subcortical Network's Energy Across Cognitive Task Conditions.** Cells indicate the median of paired differences among subjects, with p-values (in parentheses) from the Wilcoxon signed-rank test after applying multiple comparison correction. Highlighted cells represent comparisons with a corrected p-value less than 0.05.

| | Rest | 0-Back | 1-Back | 2-Back | Inhibition | Initiation | Shifting |
|---|---|---|---|---|---|---|---|
| **Rest** | - | - | - | - | - | - | - |
| **0-Back** | 0.079 (2.38E-11) | - | - | - | - | - | - |
| **1-Back** | 0.107 (1.76E-17) | 0.024 (0.088) | - | - | - | - | |
| **2-Back** | 0.115 (4.50E-17) | 0.022 (0.153) | 0.007 (0.775) | - | - | - | - |
| **Inhibition** | 0.089 (1.82E-12) | -0.011 (0.666) | -0.009 (0.017) | -0.024 (0.028) | - | - | - |
| **Initiation** | 0.081 (1.99E-12) | 0.010 (0.676) | -0.008 (0.214) | -0.026 (0.333) | 0.010 (0.353) | - | - |
| **Shifting** | 0.065 (1.41E-04) | -0.021 (7.72E-05) | -0.026 (9.30E-10) | -0.041 (3.25E-09) | -0.009 (7.91E-05) | -0.011 (3.55E-06) | - |

**Table S17. Pairwise Comparisons of Ventral Attention Network's Energy Across Cognitive Task Conditions.** Cells indicate the median of paired differences among subjects, with p-values (in parentheses) from the Wilcoxon signed-rank test after applying multiple comparison correction. Highlighted cells represent comparisons with a corrected p-value less than 0.05.

| | Rest | 0-Back | 1-Back | 2-Back | Inhibition | Initiation | Shifting |
|---|---|---|---|---|---|---|---|
| **Rest** | - | - | - | - | - | - | - |
| **0-Back** | 0.152 (9.20E-16) | - | - | - | - | - | - |
| **1-Back** | 0.156 (5.35E-19) | 0.016 (0.238) | - | - | - | - | |
| **2-Back** | 0.150 (4.00E-20) | 0.016 (0.182) | 0.004 (0.940) | - | - | - | - |
| **Inhibition** | 0.141 (1.28E-16) | -0.001 (0.940) | -0.012 (0.247) | -0.019 (0.238) | - | - | - |
| **Initiation** | 0.135 (2.32E-15) | 0.004 (0.926) | -0.005 (0.254) | -0.015 (0.247) | -0.002 (0.941) | - | - |
| **Shifting** | 0.073 (1.39E-05) | -0.076 (5.21E-08) | -0.108 (1.51E-11) | -0.088 (4.81E-13) | -0.070 (9.29E-09) | -0.079 (4.96E-08) | - |

**Table S18. Pairwise Comparisons of Visual Network's Energy Across Cognitive Task Conditions.** Cells indicate the median of paired differences among subjects, with p-values (in parentheses) from the Wilcoxon signed-rank test after applying multiple comparison correction. Highlighted cells represent comparisons with a corrected p-value less than 0.05.

| | AUD | COP | DAN | DMN | FPC | SAL | SM | SC | VAN | VIS |
|---|---|---|---|---|---|---|---|---|---|---|
| **AUD** | - | - | - | - | - | - | - | - | - | - |
| **COP** | 0.042 (0.011) | - | - | - | - | - | - | - | - | - |
| **DAN** | 0.011 (0.601) | -0.023 (0.059) | - | - | - | - | - | - | - | - |
| **DMN** | -0.037 (0.062) | -0.084 (1.27E-07) | -0.053 (0.016) | - | - | - | - | - | - | - |
| **FPC** | -0.067 (1.53E-04) | -0.117 (3.35E-11) | -0.090 (3.37E-05) | -0.036 (0.007) | - | - | - | - | - | - |
| **SAL** | -0.027 (0.116) | -0.065 (5.84E-06) | -0.039 (0.030) | -0.004 (0.999) | 0.040 (0.034) | - | - | - | - | - |
| **SM** | 0.044 (0.780) | -0.006 (0.002) | 0.028 (0.393) | 0.065 (0.063) | 0.097 (2.94E-04) | 0.073 (0.166) | - | - | - | - |
| **SC** | -0.011 (0.169) | -0.048 (0.523) | -0.021 (0.304) | 0.029 (0.003) | 0.072 (7.83E-07) | 0.017 (0.003) | -0.042 (0.079) | - | - | - |
| **VAN** | -0.012 (0.448) | -0.066 (0.001) | -0.037 (0.258) | 0.029 (0.425) | 0.052 (0.006) | 0.031 (0.448) | -0.052 (0.609) | 0.0048 (0.031) | - | - |
| **VIS** | 0.034 (0.166) | -0.008 (0.559) | 0.012 (0.321) | 0.078 (4.02E-04) | 0.124 (7.20E-07) | 0.070 (0.001) | -0.011 (0.051) | 0.045 (0.765) | 0.039 (0.036) | - |

**Table S19. Pairwise Comparisons of Networks' Energy Alterations in Transition from Rest to 0-Back Task.** Cells indicate the median of paired differences among subjects, with p-values (in parentheses) from the Wilcoxon signed-rank test after applying multiple comparison correction. Highlighted cells represent comparisons with a corrected p-value less than 0.05.

| | AUD | COP | DAN | DMN | FPC | SAL | SM | SC | VAN | VIS |
|---|---|---|---|---|---|---|---|---|---|---|
| **AUD** | - | - | - | - | - | - | - | - | - | - |
| **COP** | 0.156 (2.44E-23) | - | - | - | - | - | - | - | - | - |
| **DAN** | 0.105 (7.39E-11) | -0.036 (0.004) | - | - | - | - | - | - | - | - |
| **DMN** | 0.062 (1.41E-09) | -0.085 (4.11E-10) | -0.049 (0.024) | - | - | - | - | - | - | - |
| **FPC** | 0.016 (0.155) | -0.123 (9.63E-15) | -0.079 (2.82E-06) | -0.043 (6.03E-04) | - | - | - | - | - | - |
| **SAL** | 0.062 (2.82E-06) | -0.093 (2.65E-09) | -0.059 (0.012) | -0.008 (0.590) | 0.036 (0.007) | - | - | - | - | - |
| **SM** | 0.112 (1.45E-14) | -0.001 (1.19E-04) | 0.037 (0.820) | 0.055 (0.005) | 0.110 (1.29E-07) | 0.061 (0.005) | - | - | - | - |
| **SC** | 0.099 (1.92E-11) | -0.064 (0.141) | -0.012 (0.346) | 0.034 (0.007) | 0.076 (1.29E-07) | 0.041 (0.002) | -0.022 (0.267) | - | - | - |
| **VAN** | 0.072 (5.33E-07) | -0.060 (2.00E-05) | -0.034 (0.234) | 0.024 (0.488) | 0.066 (0.001) | 0.023 (0.306) | -0.049 (0.243) | -0.024 (0.032) | - | - |
| **VIS** | 0.106 (1.85E-11) | -0.009 (0.103) | 0.023 (0.381) | 0.075 (0.003) | 0.109 (2.87E-07) | 0.078 (0.001) | 0.002 (0.234) | 0.034 (0.727) | 0.042 (0.039) | - |

**Table S20. Pairwise Comparisons of Networks' Energy Alterations in Transition from Rest to 1-Back Task.** Cells indicate the median of paired differences among subjects, with p-values (in parentheses) from the Wilcoxon signed-rank test after applying multiple comparison correction. Highlighted cells represent comparisons with a corrected p-value less than 0.05.

| | AUD | COP | DAN | DMN | FPC | SAL | SM | SC | VAN | VIS |
|---|---|---|---|---|---|---|---|---|---|---|
| **AUD** | - | - | - | - | - | - | - | - | - | - |
| **COP** | 0.117<br>1.17E-09) | - | - | - | - | - | - | - | - | - |
| **DAN** | 0.051<br>(0.006) | -0.045<br>(0.004) | - | - | - | - | - | - | - | - |
| **DMN** | 0.016<br>(0.088) | -0.085<br>(1.29E-08) | -0.047<br>(0.124) | - | - | - | - | - | - | - |
| **FPC** | -0.031<br>(0.044) | -0.145<br>(2.94E-15) | -0.085<br>(5.54E-06) | -0.044<br>(3.55E-05) | - | - | - | - | - | - |
| **SAL** | 0.013<br>(0.116) | -0.083<br>(2.03E-08) | -0.039<br>(0.115) | -0.001<br>(0.884) | 0.052<br>(1.66E-04) | - | - | - | - | - |
| **SM** | 0.084<br>(0.016) | -0.027<br>(2.66E-05) | 0.036<br>(0.508) | 0.050<br>(0.203) | 0.104<br>(8.17E-06) | 0.073<br>(0.267) | - | - | - | - |
| **SC** | 0.036<br>(9.03E-05) | -0.068<br>(0.203) | -0.009<br>(0.172) | 0.015<br>(0.004) | 0.067<br>(1.68E-08) | 0.032<br>(0.004) | -0.020<br>(0.053) | - | - | - |
| **VAN** | 0.030<br>(0.142) | -0.068<br>(1.02E-04) | -0.030<br>(0.294) | 0.011<br>(0.656) | 0.067<br>(0.001) | 0.002<br>(0.655) | -0.043<br>(0.655) | -0.014<br>(0.023) | - | - |
| **VIS** | 0.080<br>(8.25E-05) | -0.024<br>(0.041) | 0.021<br>(0.339) | 0.057<br>(0.002) | 0.117<br>(1.16E-08) | 0.059<br>(0.003) | -0.022<br>(0.073) | 0.039<br>(0.655) | 0.032<br>(0.053) | - |

**Table S21. Pairwise Comparisons of Networks' Energy Alterations in Transition from Rest to 2-Back Task.** Cells indicate the median of paired differences among subjects, with p-values (in parentheses) from the Wilcoxon signed-rank test after applying multiple comparison correction. Highlighted cells represent comparisons with a corrected p-value less than 0.05.

| | AUD | COP | DAN | DMN | FPC | SAL | SM | SC | VAN | VIS |
|---|---|---|---|---|---|---|---|---|---|---|
| **AUD** | - | - | - | - | - | - | - | - | - | - |
| **COP** | 0.036<br>(0.065) | - | - | - | - | - | - | - | - | - |
| **DAN** | 0.017<br>(0.912) | -0.019<br>(0.159) | - | - | - | - | - | - | - | - |
| **DMN** | -0.039<br>(0.009) | -0.076<br>(3.83E-06) | -0.042<br>(0.021) | - | - | - | - | - | - | - |
| **FPC** | -0.041<br>(0.002) | -0.097<br>(3.83E-06) | -0.059<br>(0.003) | -0.006<br>(0.289) | - | - | - | - | - | - |
| **SAL** | -0.037<br>(0.037) | -0.067<br>(7.32E-05) | -0.039<br>(0.045) | 0.003<br>(0.890) | 0.016<br>(0.304) | - | - | - | - | - |
| **SM** | 0.023<br>(0.306) | -0.004<br>(0.003) | 0.013<br>(0.299) | 0.034<br>(0.217) | 0.056<br>(0.059) | 0.046<br>(0.304) | - | - | - | - |
| **SC** | -0.013<br>(0.632) | -0.046<br>(0.507) | -0.021<br>(0.686) | 0.026<br>(0.027) | 0.039<br>(0.004) | 0.017<br>(0.037) | -0.025<br>(0.162) | - | - | - |
| **VAN** | -0.022<br>(0.178) | -0.062<br>(0.003) | -0.023<br>(0.191) | 0.009<br>(0.599) | 0.015<br>(0.268) | 0.012<br>(0.686) | -0.059<br>(0.706) | -0.004<br>(0.095) | - | - |
| **VIS** | 0.028<br>(0.432) | -0.020<br>(0.412) | 0.005<br>(0.545) | 0.058<br>(0.001) | 0.052<br>(2.08E-04) | 0.043<br>(0.003) | 0.003<br>(0.068) | 0.051<br>(0.891) | 0.032<br>(0.046) | - |

**Table S22. Pairwise Comparisons of Networks' Energy Alterations in Transition from Rest to Inhibition Condition of Go/No-go Task.** Cells indicate the median of paired differences among subjects, with p-values (in parentheses) from the Wilcoxon signed-rank test after applying multiple comparison correction. Highlighted cells represent comparisons with a corrected p-value less than 0.05.

| | AUD | COP | DAN | DMN | FPC | SAL | SM | SC | VAN | VIS |
|---|---|---|---|---|---|---|---|---|---|---|
| **AUD** | - | - | - | - | - | - | - | - | - | - |
| **COP** | 0.037 (0.026) | - | - | - | - | - | - | - | - | - |
| **DAN** | -0.002 (0.647) | -0.036 (0.013) | - | - | - | - | - | - | - | - |
| **DMN** | -0.042 (0.002) | -0.086 (1.67E-09) | -0.038 (0.034) | - | - | - | - | - | - | - |
| **FPC** | -0.050 (0.002) | -0.086 (6.98E-09) | -0.047 (0.018) | 0.002 (0.559) | - | - | - | - | - | - |
| **SAL** | -0.034 (0.009) | -0.076 (1.16E-07) | -0.034 (0.057) | 0.001 (0.884) | 0.010 (0.631) | - | - | - | - | - |
| **SM** | 0.024 (0.224) | -0.005 (7.87E-05) | 0.034 (0.612) | 0.048 (0.022) | 0.060 (0.010) | 0.076 (0.081) | - | - | - | - |
| **SC** | -0.018 (0.559) | -0.050 (0.221) | -0.003 (0.315) | 0.041 (0.004) | 0.036 (0.002) | 0.029 (0.006) | -0.01519 (0.171) | - | - | - |
| **VAN** | -0.014 (0.232) | -0.058 (8.42E-04) | -0.026 (0.512) | 0.022 (0.141) | 0.033 (0.081) | 0.027 (0.221) | -0.050 (0.754) | -0.001 (0.097) | - | - |
| **VIS** | 0.031 (0.171) | -0.011 (0.483) | 0.033 (0.084) | 0.056 (1.23E-06) | 0.064 (1.23E-06) | 0.067 (1.21E-05) | -0.001 (0.005) | 0.049 (0.572) | 0.058 (0.010) | - |

**Table S23. Pairwise Comparisons of Networks' Energy Alterations in Transition from Rest to Initiation Condition of Go/No-go Task.** Cells indicate the median of paired differences among subjects, with p-values (in parentheses) from the Wilcoxon signed-rank test after applying multiple comparison correction. Highlighted cells represent comparisons with a corrected p-value less than 0.05.

| | AUD | COP | DAN | DMN | FPC | SAL | SM | SC | VAN | VIS |
|---|---|---|---|---|---|---|---|---|---|---|
| **AUD** | - | - | - | - | - | - | - | - | - | - |
| **COP** | 0.033 (0.018) | - | - | - | - | - | - | - | - | - |
| **DAN** | -0.045 (0.002) | -0.077 (1.68E-06) | - | - | - | - | - | - | - | - |
| **DMN** | -0.008 (0.800) | -0.036 (0.002) | 0.044 (0.002) | - | - | - | - | - | - | - |
| **FPC** | -0.055 (4.49E-06) | -0.101 (1.64E-11) | -0.014 (0.434) | -0.05 (1.05E-06) | - | - | - | - | - | - |
| **SAL** | -0.050 (0.025) | -0.069 (2.36E-06) | 0.017 (0.314) | -0.025 (0.025) | 0.024 (0.017) | - | - | - | - | - |
| **SM** | 0.050 (0.063) | 0.001 (3.85E-06) | 0.081 (0.114) | 0.022 (0.062) | 0.112 (0.001) | 0.049 (0.650) | - | - | - | - |
| **SC** | -0.039 (0.118) | -0.067 (0.800) | 0.025 (2.42E-05) | -0.022 (0.063) | 0.039 (3.59E-08) | -0.005 (3.99E-04) | -0.047 (0.001) | - | - | - |
| **VAN** | -0.002 (0.800) | -0.031 (0.018) | 0.033 (0.007) | -0.001 (0.982) | 0.061 (1.93E-05) | 0.031 (0.051) | -0.050 (0.098) | 0.021 (0.096) | - | - |
| **VIS** | -0.017 (0.672) | -0.035 (0.007) | 0.040 (0.029) | -0.006 (0.800) | 0.057 (3.79E-04) | 0.026 (0.098) | -0.048 (0.201) | 0.025 (0.033) | -0.001 (0.773) | - |

**Table S24. Pairwise Comparisons of Networks' Energy Alterations in Transition from Rest to Shifting Task.** Cells indicate the median of paired differences among subjects, with p-values (in parentheses) from the Wilcoxon signed-rank test after applying multiple comparison correction. Highlighted cells represent comparisons with a corrected p-value less than 0.05.

| Input Features | Accuracy | Kappa | Per Class Accuracy | | | | |
|---|---|---|---|---|---|---|---|
| | | | Rest | Working Memory | Inhibitory Control | Cognitive Flexibility | Balanced Accuracy |
| Network Energy | 0.577 | 0.24 | 0.601 | 0.78 | 0.339 | 0.367 | 0.527 |
| Global Clustering Coefficient | 0.464 | 0.135 | 0.218 | 0.82 | 0.276 | 0.015 | 0.332 |
| Global Efficiency | 0.52 | 0.268 | 0.52 | 0.802 | 0.205 | 0.3 | 0.457 |
| Global Modularity | 0.481 | 0.166 | 0.386 | 0.865 | 0.17 | 0.043 | 0.366 |
| All Global Network Measures | 0.61 | 0.415 | 0.697 | 0.794 | 0.43 | 0.33 | 0.563 |

**Table S25. Performance of Cognitive Task Classification for Each Set of Input Features**

| Input Features | $R^2$ | MAE |
| --- | --- | --- |
| Network Energy | 0.447 | 11.294 |
| Global Clustering Coefficient | 0.439 | 12.795 |
| Global Efficiency | 0.277 | 13.249 |
| Global Modularity | 0.394 | 12.442 |
| All Global Network Measures | 0.543 | 10.36 |

**Table S26. Performance of Predicting Chronical Age for Each Set of Input Features.**

| Task | P-value | Median of Differences |
| --- | --- | --- |
| Resting | << .001 | -4.05E-2 |
| 0-Back | << .001 | -3.37E-2 |
| 1-Back | << .001 | -3.39E-2 |
| 2-Back | << .001 | -3.17E-2 |
| Inhibition | << .001 | -2.51E-2 |
| Initiation | << .001 | -3.52E-2 |
| Shifting | << .001 | -2.59E-2 |

**Table S27. Pairwise Comparison of Energy between Whole-brain Networks during Cognitive Task Conditions and Their Corresponding Random Topology Networks**

| Task | ICC | Lower Band Confidence Interval | Upper Band Confidence Interval | P-value |
| --- | --- | --- | --- | --- |
| Resting | 0.94250563 | 0.9200982 | 0.95862918 | 2.89E-51 |
| 0-Back | 0.915003 | 0.8818769 | 0.93883931 | 1.07E-40 |
| 1-Back | 0.90190533 | 0.86367464 | 0.92941471 | 5.56E-37 |
| 2-Back | 0.90647835 | 0.87002993 | 0.93270529 | 3.29E-38 |
| Inhibition | 0.92396689 | 0.89433431 | 0.94528939 | 1.19E-43 |
| Initiation | 0.89641186 | 0.8560402 | 0.92546182 | 1.35E-35 |
| Shifting | 0.93578342 | 0.91075613 | 0.95379213 | 3.21E-48 |

**Table S28. Intraclass Correlation Statistics between Whole-brain Energy of Power's Atlas-based Networks and Schaefer's Atlas-based Networks.**
